# Impaired phagocytic function in CX3CR1^+^ tissue-resident skeletal muscle macrophages prevents muscle recovery after influenza A virus-induced pneumonia in aged mice

**DOI:** 10.1101/833236

**Authors:** Constance E. Runyan, Lynn C. Welch, Emilia Lecuona, Masahiko Shigemura, Luciano Amarelle, Hiam Abdala-Valencia, Nikita Joshi, Ziyan Lu, Kiwon Nam, Nikolay S. Markov, Alexandra C. McQuattie-Pimentel, Raul Piseaux-Aillon, Yuliya Politanska, Lango Sichizya, Satoshi Watanabe, Kinola J.N. Williams, GR Scott Budinger, Jacob I. Sznajder, Alexander V. Misharin

**Author notes:** equal contribution (Jacob I. Sznajder, Alexander V. Misharin).

## Abstract

Skeletal muscle dysfunction in survivors of pneumonia is a major cause of lasting morbidity that disproportionately affects older individuals. We found that skeletal muscle recovery was impaired in aged compared with young mice after influenza A virus-induced pneumonia. In young mice, recovery of muscle loss was associated with expansion of tissue-resident skeletal muscle macrophages and downregulation of MHC II expression, followed by a proliferation of muscle satellite cells. These findings were absent in aged mice and in mice deficient in *Cx3cr1*. Transcriptomic profiling of tissue-resident skeletal muscle macrophages from aged compared with young mice showed downregulation of pathways associated with phagocytosis and proteostasis, and persistent upregulation of inflammatory pathways. Consistently, skeletal muscle macrophages from aged mice failed to downregulate MHCII expression during recovery from influenza A virus induced pneumonia and showed impaired phagocytic function *in vitro*. Like aged animals, mice deficient in the phagocytic receptor *Mertk* showed no macrophage expansion, MHCII downregulation or satellite cell proliferation and failed to recover skeletal muscle function after influenza A pneumonia. Our data suggest that a loss of phagocytic function in a CX3CR1^+^ tissue-resident skeletal muscle macrophage population in aged mice precludes satellite cell proliferation and recovery of skeletal muscle function after influenza A pneumonia.

## INTRODUCTION

While elderly individuals are at increased risk of developing and dying from pneumonia, most patients with access to modern health care survive (Jain et al., 2015; Thannickal et al., 2015). A growing body of literature suggests that elderly pneumonia survivors are at increased risk of developing age-related disorders including persistent lung dysfunction (Mittl et al., 1994), skeletal muscle atrophy limiting mobility (Herridge et al., 2003), myocardial infarction (Corrales-Medina Vicente et al., 2012) chronic kidney disease (Murugan et al., 2010), dementia (Tate et al., 2014) and cognitive impairment (Girard et al., 2018). Skeletal muscle dysfunction develops in the majority of survivors of pneumonia and may persist for years after hospital discharge, where it is a major driver of morbidity (Chan et al., 2018; Dos Santos et al., 2016; Herridge et al., 2003; Herridge et al., 2011; Walsh et al., 2014). Persistent skeletal muscle weakness in survivors of severe pneumonia disproportionately affects elderly patients, resulting in reduced quality of life including an increased risk of hospitalizations, long-term disability, and loss of independence (Barreiro et al., 2015; Falsey et al., 2005; Gozalo et al., 2012; Herridge et al., 2011; Pfoh et al., 2016).

Because skeletal muscle fibers are post-mitotic, muscle myofibers lost during injury must be replaced through activation, proliferation, and differentiation of muscle satellite cells, a muscle-specific progenitor cell population (Brack and Rando, 2012; Wang and Rudnicki, 2011). After injury, muscle satellite cells undergo rapid proliferation, expand in number and fuse with damaged myofibers, restoring the number of myonuclei, myofiber content and volume (Munoz-Canoves and Michele, 2013; Sala et al., 2019; Tierney et al., 2014). The process of muscle repair has been studied in models of direct injury, for example freezing, trauma, and toxin injection, but the mechanisms by which muscle fibers are regenerated after the indirect injury that develops after pneumonia are not known.

Infection with the influenza A virus is among the most common causes of pneumonia and the morbidity and mortality attributable to influenza A pneumonia disproportionately impacts the elderly (Jain et al., 2015; Ortiz et al., 2013). Influenza A virus-induced pneumonia in mice is an attractive model to study the effects of aging on pneumonia-induced muscle function as it recapitulates worsened skeletal muscle dysfunction reported in older survivors of pneumonia (Bartley et al., 2016). Furthermore, as the influenza A virus is trophic for the lung epithelium, there is little if any viremia (Radigan et al., 2019). Therefore, the muscle loss after influenza A pneumonia is likely attributable to endocrine factors common to pneumonia secondary to a variety of pathogens, rather than the virus itself. Here we report that when the dose of influenza A administered to young and aged mice is titrated to induce nearly equivalent mortality, surviving young but not aged mice recovered muscle function after pneumonia. In young mice, a population of *Cx3cr1*-expressing tissue-resident macrophages in the skeletal muscle expanded and downregulated MHCII expression after influenza A infection in the absence of recruitment of monocytes from the bone marrow. This was followed by proliferation of muscle satellite cells. All of these findings were absent in aged mice and mice deficient in *Cx3cr1*. Transcriptomic profiling suggested a loss of phagocytic function in tissue-resident macrophages from the skeletal muscle of aged mice, which was confirmed using *in vitro* assays. Knockout of *Mertk*, a tyrosine kinase involved in macrophage phagocytosis, phenocopied the molecular and physiologic findings in aged mice. Our findings implicate phagocytosis-induced signaling in *Cx3cr1*-expressing, tissue-resident skeletal muscle macrophages driving satellite cell proliferation during muscle recovery after pneumonia-induced atrophy. These findings may have broad implications for other systemic inflammatory conditions associated with age-related muscle dysfunction.

## RESULTS

### Aged mice do not recover skeletal muscle function after influenza A-induced pneumonia

When infected with the same dose of influenza A virus, aged mice exhibit increased mortality (Figure S1A). Therefore, to examine muscle recovery after injury, we titrated the dose of influenza A virus in young (4 months old) and aged (20-24 months old) mice to achieve ∼25% mortality and similar levels of weight loss in each group (Figure S1B). We continuously measured voluntary running distance on monitored exercise wheels in young and aged mice over the course of influenza A infection and recovery, as a measure of muscle function. Mice adapted to the wheels over 14 days, after which the voluntary distance run on the wheel was consistent and similar between animals (Figure 1A), although on average it was lower in aged mice (Figure S2A). Once baseline running distances were established, mice were infected with influenza A virus, and then monitored continuously for up to 60 days. Both young and old mice almost stopped running after the influenza A virus infection, reaching a minimum at day 10 post infection (Figure 1A, B). However, by day 20, the distance run returned to pre-infection levels in young mice. In contrast, old mice failed to return to their pre-infection activity levels (Figure 1A, B). Similarly, while influenza A virus pneumonia induced a loss of forearm grip strength in both young and old mice (Figure S2B), aged mice failed to recover forearm grip strength 30 and 60 days after influenza A virus infection (Figure 1C).

**Figure 1.**
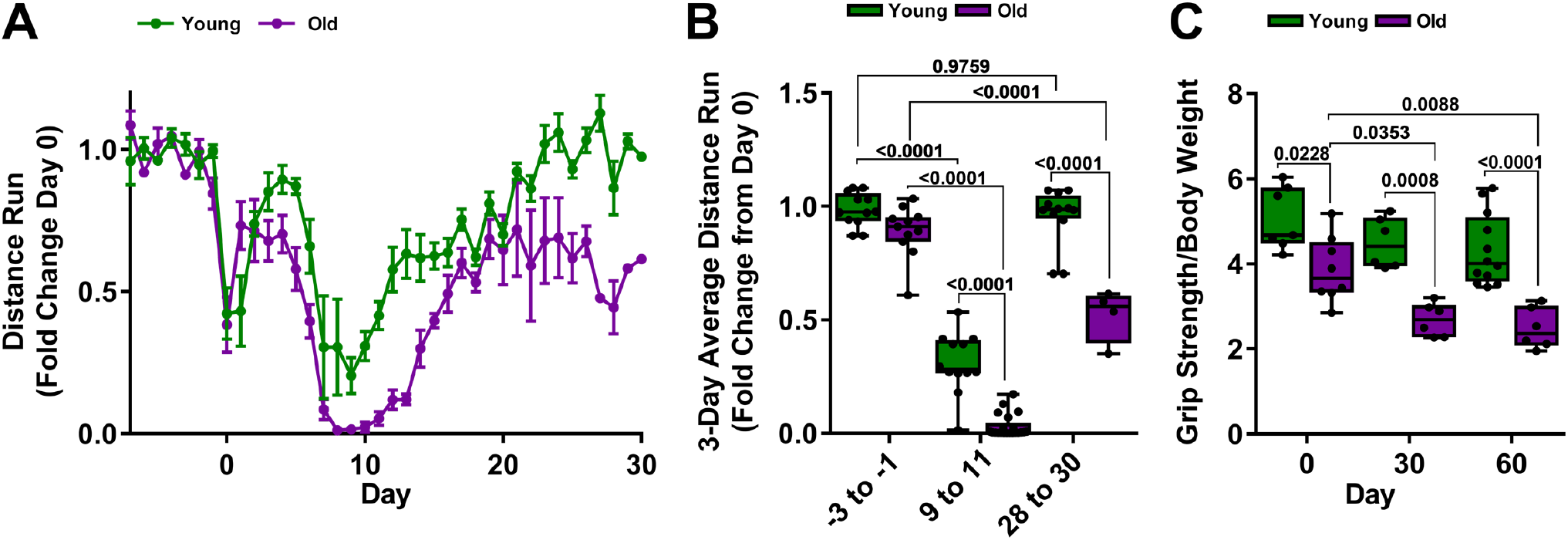
Old mice fail to recover skeletal muscle function after influenza A infection. (A) Monitored running wheels were placed in cages and the voluntary distance run was measured for 14 days before animals were infected with influenza A virus (IAV). At Day 0, young (4 months old; green lines) mice were infected with 25 pfu of IAV and aged (20-24 months old; purple lines) mice were infected with 15 pfu IAV. Voluntary wheel running was recorded for 30 days and the 8 PM to 6 AM average wheel rotation/mouse, normalized to the 10-day average control is shown (n=12 mice per group). A graph of the distance run is shown with means ± S.D. See also Figure S1 and S2. (B) Three-day average voluntary wheel running distance per cage normalized to the 10-day average control is shown for young mice (4 months old; green bars) and aged mice (20-24 months old; purple bars). The average distance run for Day −3 to −1 prior to infection with influenza A virus (IAV) (25 pfu for young mice, 15 pfu for old mice), day 9-11 post infection, and day 28 to 30 post infection is shown (n=12 mice per group). Box plot center lines are median, box limits are upper and lower quartiles, whiskers are minimal and maximal values, a two-way ANOVA with Tukey post-hoc corrections for comparison with more than three groups was used to determine statistical differences, and p values are shown on graph. Each dot represents a single animal. (C) Young (4 months old; green bars) and aged (20-24 months old; purple bars) mice were infected with IAV at Day 0 (25 pfu for young mice, 15 pfu for old mice). Forelimb grip strength was measured at the indicated times (n=6-12 mice per group). Box plot center lines are median, box limits are upper and lower quartiles, whiskers are minimal and maximal values, a two-way ANOVA with Tukey post-hoc corrections for comparison with more than three groups was used to determine statistical differences, and p values are shown on graph. Each dot represents a single animal.

Despite the markedly reduced dose of virus and similar mortality, aged mice still had worse lung injury as measured by bronchoalveolar lavage protein content 10 days after infection (Figure 2A, B) and increased levels IL-6 in bronchoalveolar lavage fluid (Figure 2C) and in the plasma compared to young mice (Figure 2D). The soleus muscle wet weight decreased similarly in both young and old mice (Figure 2E), however, the loss of wet weight in the *extensor digitorum longus* (EDL) muscle was more pronounced in aged mice (Figure 2F). Average soleus cross-sectional area was reduced in both young and aged mice after influenza A virus-induced pneumonia, but aged mice had significantly reduced fiber volume before influenza A infection (Figure 2G). Similarly, the left-shift of the distribution of the cross-sectional area measurements after influenza A virus infection was less pronounced in aged mice because of a generally lower initial muscle volume (Figure S3). We previously reported that systemic increase in IL-6 level is a direct cause of muscle atrophy after influenza A infection through induction of the atrophy-promoting E3 ubiquitin ligase, Atrogin-1 (*Fbxo32*) (Radigan et al., 2019). The maximal induction of Atrogin-1 in the skeletal muscle was similar in young and aged animals (Figure 2H,I).

**Figure 2.**
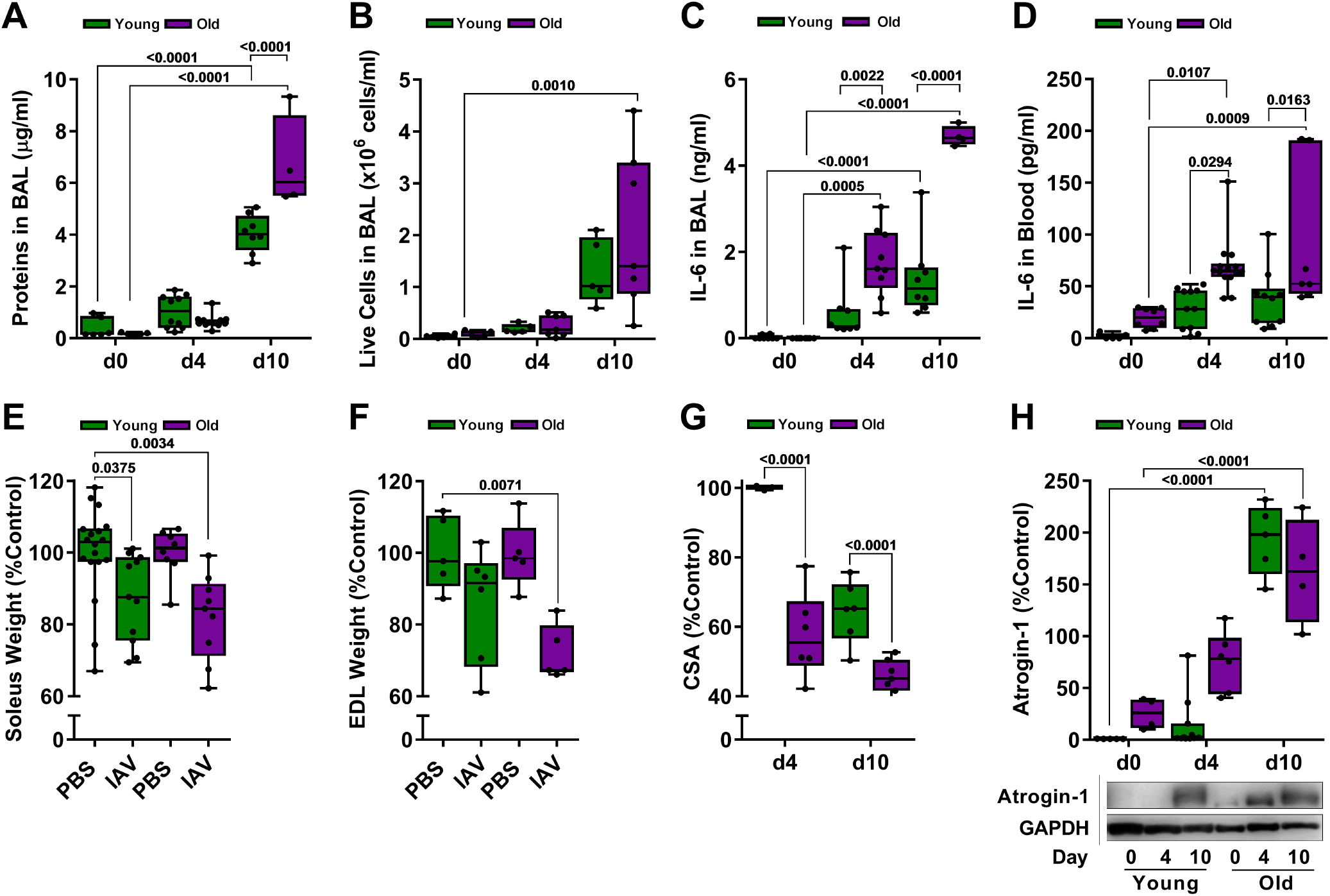
Influenza A virus infection induces muscle loss in both aged and young mice. For all data young (4 months old; green bars) and old (20-24 months old; purple bars) mice were infected with Influenza A virus (25 pfu for young mice, 15 pfu for aged mice). Mice were harvested for analysis at 0, 4 and 10 days post-infection. A two-way ANOVA with Tukey post-hoc corrections for comparison with more than three groups was used to determine statistical differences, p values are shown on graph. Box plot center lines are median, box limits are upper and lower quartiles. Each dot represents an individual animal. (A) Total protein in bronchoalveolar lavage (BAL) fluid was measured by Bradford assay (n=4-12 mice). (B) Number of live cells in the BAL were counted using hemocytometer, dead cells were excluded using trypan blue (n=5-7 mice). (C) IL-6 levels in the serum were measured by ELISA (n=4-8 mice). (D) Soleus muscles were excised and soleus wet muscle weights were determined. Values are presented as a percent of naive age-matched controls (n=8-18 mice). (E) *Extensor digitorum longus* (EDL) muscles were excised and EDL wet muscle weights were measured. Values are presented as a percent of naive age-matched controls (n=5-7 mice). (F) Soleus cross-sectional areas were determined using transverse cross-sections immunostained for laminin. Values are normalized to naive age-matched controls (n=4-7 mice). (G) Quantification (top panel) and representative Western blot (bottom panel) of Atrogin-1 expression in the mouse tibia anterior muscle. Values are normalized to GAPDH (loading control) (n=5-11 mice).

### Satellite cells and fibroadipogenic progenitors fail to expand in aged mice after influenza A virus-induced pneumonia

We used flow cytometry to simultaneously quantify muscle satellite cells (MuSC) and fibro-adipogenic progenitors (FAP) as well as lymphoid and myeloid immune cells during muscle repair. We identified MuSC as CD45^−^CD31^−^CD34^+^Sca-1^−^Itga7^+^, and FAP as CD45^−^CD31^−^CD34^+^Sca-1^+^Itga7^+^ (Figure 3A). In young, but not in aged mice, we observed an expansion of MuSC and FAP after influenza A virus-induced pneumonia (Figure 3B-G). This suggests that muscle dysfunction induced by influenza A pneumonia is sufficient to induce regenerative cell populations in young mice, but that this response is impaired with age.

**Figure 3.**
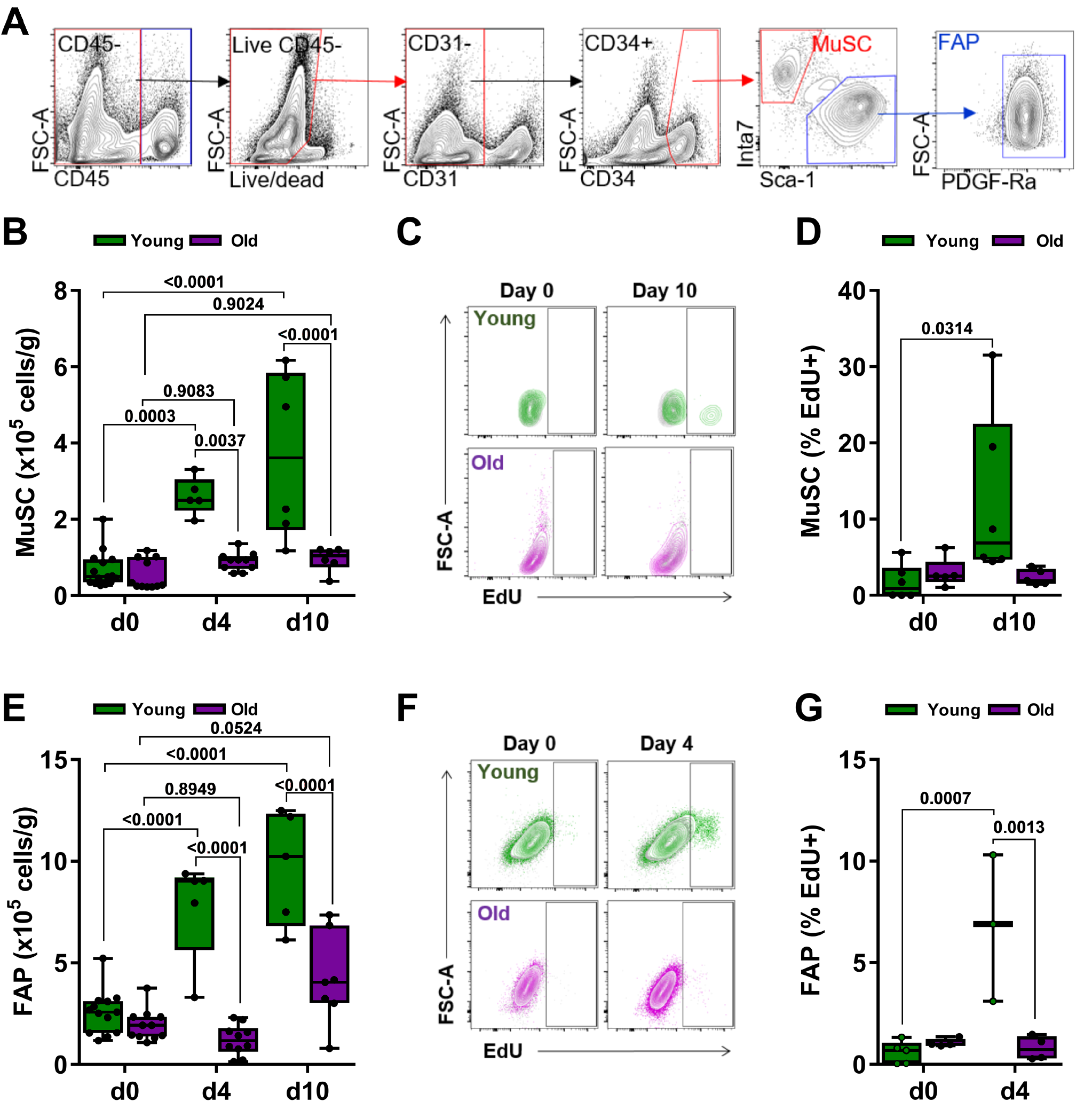
Expansion of muscle satellite cells and fibroadipogenic progenitors is reduced in aged compared with young mice after influenza A virus pneumonia. For all experiments, young (4 months old; green bars) and aged (20-24 months old; purple bars) mice were infected with Influenza A virus (25 pfu for young mice, 15 pfu for aged mice). Mice were harvested for analysis before the infection (Day 0) and 4 and 10 days after infection. A two-way ANOVA with Tukey post-hoc corrections for comparison with more than three groups was used to show statistical differences. Box plot center lines are median, box limits are upper and lower quartiles, whiskers are minimal and maximal values, p values are shown on the graph. Each dot represents an individual animal. (A) Representative gating strategy to identify muscle satellite cells (MuSC) and fibroadipogenic progenitors (FAP). (B) Enumeration of MuSC using flow cytometry. Values shown are per gram of tissue (n=6-11 mice). (C) Young (top 2 panels; green) or aged mice (bottom 2 panels; purple) were injected with 2 mg EdU, 16 hours prior to day 0 or day 10. Representative contour plots showing percentage of EdU-positive MuSC are shown. Plots from EdU-injected mice are overlaid against plots of a separate, non-EdU-injected control mouse (shown in grey). (D) Box and whiskers plot showing percent EdU+ MuSC of total MuSC at day 0, and 10 (n=5-6 mice). (E) Enumeration of FAP using flow cytometry. Values shown are per gram of tissue (n=5-13 mice). (F) Young (top 2 panels; green) or aged mice (bottom 2 panels; purple) were injected with 2 mg EdU 16 hours prior to day 0 or day 10. Representative contour plots showing percentage of EdU-positive FAP are shown. Plots from EdU-injected mice are overlaid against plots of a separate, non-EdU-injected control mouse (shown in grey). (G) Box and whiskers plot showing percent EdU+ FAP of total FAP at day 0 and day 4 (n=3-5 mice).

### Tissue-resident CX3CR1+ skeletal muscle macrophages expand and downregulate MHCII during recovery from influenza A virus-induced pneumonia

We measured myeloid and lymphoid populations in the skeletal muscle after influenza A infection using flow cytometry (Figure 4A and S4M). In young mice, we observed increases in the number of skeletal muscle macrophages (CD45+Ly6G-Siglec F-NK1.1-CD11b+Ly6C-CD64+) after influenza A infection (Figure 4B). This increase in skeletal muscle macrophages was not observed in aged mice (Figure 4B). In contrast, we did not observe influenza A virus-induced increased numbers of neutrophils, eosinophils, natural killer cells, CD11b+ dendritic cells, Ly6C-high classical monocytes or Ly6C-low non-classical monocytes, CD4 and CD8 T cells, regulatory T cells, or B cells in the skeletal muscle of either young or aged mice after influenza A infection (Figure S4).

**Figure 4.**
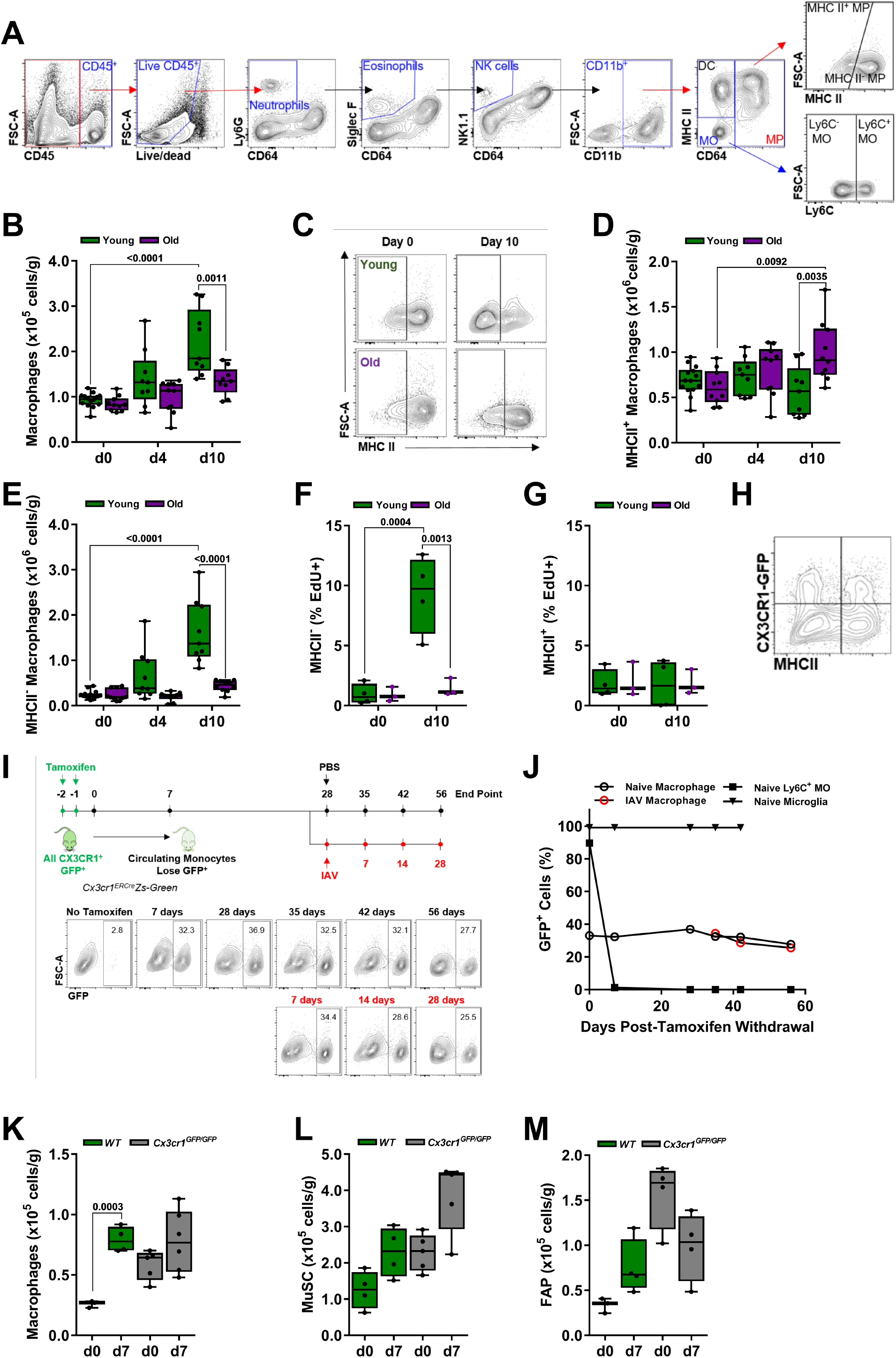
Tissue-resident skeletal muscle macrophages expand and downregulate MHCII expression after influenza A pneumonia in young, but not in aged mice. (A) Gating strategy to identify myeloid cell populations in skeletal muscle tissue using flow cytometry. (B) Young (4 months old; green bars) and aged (20-24 months old; purple bars) mice were infected with influenza A virus (25 pfu for young mice, 15 pfu for aged mice). Graph shows number of macrophages per gram of tissue, before infection (Day 0) and 4 and 10 post-infection (n=9-14 mice). (C) Representative flow cytometry showing MHCII staining in young (top 2 panels) or aged (bottom 2 panels) muscle macrophages before and 10 days after IAV infection. (D) Young (4 months old; green bars) and aged (20-24 months old; purple bars) mice were infected with influenza A virus (25 pfu for young mice, 15 pfu for aged mice). Graph shows number of MHCII^high^ macrophages in skeletal muscle per gram of tissue before infection (Day 0) and 4 and 10 days after influenza A infection (n=9-14 mice). (E) Young (4 months old; green bars) and aged (20-24 months old; purple bars) mice were infected with influenza A virus (25 pfu for young mice, 15 pfu for aged mice). Graph shows number of MHCII^low^ macrophages in skeletal muscle per gram of tissue before infection (Day 0) and 4 and 10 days after influenza A infection (n=9-14 mice). (F) Young (4 months old; green bars) and aged (20-24 months old; purple bars) mice were infected with influenza A virus (25 pfu for young mice, 15 pfu for aged mice). Graph shows percentage of EdU^+^ MHCII^high^ macrophages, of total MHCII^high^ macrophages, in skeletal muscle before infection (Day 0) and 10 days after influenza A infection (n=3-4 mice). (G) Young (4 months old; green bars) and aged (20-24 months old; purple bars) mice were infected with influenza A virus (25 pfu for young mice, 15 pfu for aged mice). Graph shows percentage of EdU^+^ MHCII^low^ macrophages, of total MHCII^low^ macrophages, in skeletal muscle before infection (Day 0) and 10 days after influenza A infection (n=3-4 mice). (H) Representative plot of macrophages from *Cx3cr1^Gfp/WT^* mice showing heterogeneous expression of MHCII and CX3CR1 (GFP). (I) *Cx3cr1^ERCre^/zsGreen* mice were administered tamoxifen (10 mg via oral gavage, twice, 24 hours apart). Mice were harvested 7, 28, 35, 42 and 56 days after the tamoxifen pulse. A separate cohort of mice was infected with 25 pfu IAV 28 days after the tamoxifen pulse and harvested 7, 14 and 28 days later. Representative flow cytometry plots of macrophage GFP expression at each of these time points are shown. (J) Refers to the schematic in (I). *Cx3cr1^ERCre^/zsGreen* mice were administered a pulse of tamoxifen (10 mg via oral gavage, twice, 24 hours apart) at Day 0. The percentage of GFP^+^ cells from the indicated cell populations is shown. Some mice were infected with 25 pfu IAV 28 days after the tamoxifen pulse and harvested 7, 14 and 28 days later. Black symbols indicate naïve mice. Red symbols indicate influenza A virus infected mice. (K) Young wild type mice (4 months old; green bars) and age-matched *Cx3cr1^Gfp/Gfp^*mice (grey bars) were infected with 25 pfu of IAV. Graph represents skeletal muscle macrophage number before infection (Day 0) and 7 days after influenza A infection. Values are expressed per gram of tissue (n=4-5 mice). (L) Young wild type mice (4 months old; green bars) and age-matched *Cx3cr1^Gfp/Gfp^* mice (grey bars) were infected with 25 pfu of IAV. Graph represents muscle satellite cell (MuSC) number before infection (Day 0) and 7 days after influenza A infection. Values are expressed per gram of tissue (n=4-5 mice). (M) Young wild type mice (4 months old; green bars) and age-matched *Cx3cr1^Gfp/Gfp^* mice (grey bars) were infected with 25 pfu of IAV. Graph represents fibroadipogenic progenitors (FAP) number before infection (Day 0) and 7 days after influenza A infection. Values are expressed per gram of tissue (n=4-5 mice). For all experiments, a two-way ANOVA with Tukey post-hoc corrections for comparison with more than three groups was used to show statistical differences, p values are shown on the graph. Each dot represents an individual animal. Also see Figure S4.

As we only observed a significant expansion in macrophages after influenza A infection in young but not aged mice, we focused our attention on these cells. In naïve mice, the majority of macrophages were represented by an MHCII^high^ population (Figure 4C). During recovery from influenza A infection, the majority of macrophages were MHCII^low^ in young, but not in aged mice (Figure 4C-E). Furthermore, incorporation of EdU into MHCII^low^ macrophages in response to pneumonia-induced muscle atrophy increased in young but not aged mice, indicating an age-dependent proliferation specific to this population (Figure 4F,G). Using reporter mice in which GFP is knocked into the Cx3cr1 gene (*Cx3cr1^GFP/+^*), we found that approximately 25-30% of skeletal muscle macrophages expressed *Cx3cr1* in naïve mice (Figure S5), and both CX3CR1^+^ and CX3CR1^-^ macrophages included populations that were MHCII^high^ and MHCII^low^ (Figure 4H). To determine whether these skeletal muscle macrophage populations are maintained by local proliferation or are replenished by monocytes recruited from the circulation, we performed fate-mapping experiments using *Cx3cr1^ER-Cre^*mice crossed to a *ZsGreen^LSL^* reporter. We administered a single pulse of tamoxifen via oral gavage to *Cx3cr1^ER-Cre^/ZsGreen^LSL^* mice to permanently label all *Cx3cr1*-expressing cells with GFP, and analyzed number of GFP-positive cells in several myeloid populations over the course of 56 days (Figure 4I,J). Microglia, which are tissue-resident brain macrophages that are not replaced by circulating monocytes over the lifespan of mice and are characterized by constitutive expression of *Cx3cr1* (Yona et al., 2013), were used as a positive control. Nearly 100% of microglia were labeled, and remained GFP-positive after the tamoxifen pulse over the course of the experiment (Figure 4J). Seven days after the tamoxifen pulse, 80% of Ly6C+ classical monocytes in the blood were GFP positive (Figure 4J). Consistent with the short half-life of circulating monocytes, almost all of GFP-positive monocytes were replaced by GFP-negative monocytes 28 days after the tamoxifen pulse (Figure 4J) (Yona et al., 2013). Similarly, 28 days after the tamoxifen pulse nearly all Ly6C-non-classical monocytes, and CD11b+ dendritic cells were also replaced by GFP-negative cells (not shown). Consistent with our findings using *Cx3cr1^Gfp/WT^* mice, we found that ∼30% of skeletal muscle macrophages were GFP-positive after the tamoxifen pulse (Figure 4J). The number of GFP-positive skeletal muscle macrophages remained stable 56 days after the tamoxifen pulse. As such, these cells represent a *bona fide,* long-lived tissue-resident macrophage population that do not require circulating monocytes to maintain their population with a half-life of at least several months.

We used this system to determine whether the increase in skeletal muscle macrophages we observed after influenza A virus-induced pneumonia was attributable to the recruitment of monocyte-derived macrophages. We treated *Cx3cr1^ER-Cre^/ZsGreen^LSL^*with a pulse of tamoxifen, waited 28 days for GFP-labeled monocytes todisappear, infected the mice with influenza A virus and then harvested mice 7, 14 and 28 days after infection (Figure 4I). We did not observe any reduction in the percentage of GFP-positive macrophages, excluding the possibility that increase in skeletal muscle macrophages following influenza A infection represented an influx of circulating cells (Figure 4I and J). Further, the finding that the number of macrophages expanded after influenza A infection combined with the constant percentage of GFP+ cells suggests both CX3CR1-positive and CX3CR1-negative cells expand proportionately after influenza A infection (Figure 4B and J). CX3CR1 is required for the adhesion and migration of leukocytes, and in the microglia CX3CR1 is necessary for synaptic pruning during development (Paolicelli et al., 2011). We therefore sought to determine whether CX3CR1 is necessary for the expansion of macrophages and recovery of muscle function after influenza A infection. We treated mice deficient in *Cx3cr1* (*Cx3cr1^Gfp/Gfp^*) with influenza A virus and performed flow cytometry on skeletal muscle during recovery. Mice deficient in *Cx3cr1* showed increased numbers of skeletal muscle macrophages at baseline, perhaps reflecting a failure to populate the niche. However, *Cx3cr1^Gfp/Gfp^* mice showed no expansion of skeletal muscle macrophages, no change in the proportion of MHCII^low^ macrophages, and no proliferation of MuSC or FAP after influenza A virus-induced pneumonia (Figure 4K-M).

### Transcriptomic profiling identifies downregulation of phagocytic and proteostatic pathways and upregulation of inflammatory pathways in tissue-resident skeletal muscle macrophages from aged compared with young mice

To identify possible mechanisms responsible for the lack of a reparative response in aged tissue-resident skeletal macrophages during recovery from influenza A virus-induced pneumonia, we performed transcriptomic profiling (RNA-seq) of flow cytometry-sorted skeletal muscle macrophages. Out of 1407 differentially expressed genes (FDR q < 0.05), 582 were upregulated in aged compared with young animals and 825 were downregulated (Figure 5A). Macrophages from naïve aged animals demonstrated decreased expression of genes involved lipid storage and metabolism (*Ldlrap1, Lrp1, Fabp4, Nr1h3*), macrophage activation and cytokine production (*Nlrp3*, *Sphk1, Msr1, Il10*), and extracellular matrix remodeling (*Spp1, Mmp13, Mmp14, Thbs1, Chil3*). (Figure 5A and Supplementary table S1 and S2, list of DEGs and corresponding GO processes). In addition, macrophages from aged animals demonstrated decreased expression of the genes involved in response to unfolded protein and cellular stress, particularly molecular chaperones (*Dnaja2, Dnajb4, Dnajb11, Hspa9, Hsp90b1, Hspbap1*), as well as several molecules involved in positive regulation of phagocytosis (*C5ar1, Msr1*). In contrast, macrophages from young naïve animals were characterized by decreased expression of the genes involved in negative regulation of cell migration (*Sema4a, Sema6d, Plxna3, Sema4c, Sema6a*) and antigen processing and presentation (*Cd74, H2-Ab1, H2-Eb1, H2-Aa*). We also performed transcriptomic profiling of flow cytometry sorted skeletal muscle macrophages from young and aged mice 60 days after IAV pneumonia. Of 1054 differentially expressed genes (FDR q < 0.05) 673 genes were upregulated in skeletal muscle macrophages from aged mice and were enriched for the genes involved in inflammation (*Il6, Tnf, Il10, Il1a, Il1b, Tnfsf14*) and proteostatic stress (*Hspa9, Hspa5, Hspb8, Hspbap1, Dnaja4, Dnajb4, Dnaja2*) (Figure 5B and Supplementary table S3 and S4, list of DEGs and corresponding GO processes). These transcriptomic data suggested impaired phagocytosis and increased inflammatory response in skeletal muscle macrophages from old mice. Consistent with these observations, we found that flow cytometry sorted tissue-resident skeletal muscle macrophages from old mice exhibited reduced uptake of fluorescently labeled polystyrene beads when compared with those from young animals (Figure 5C, D).

**Figure 5.**
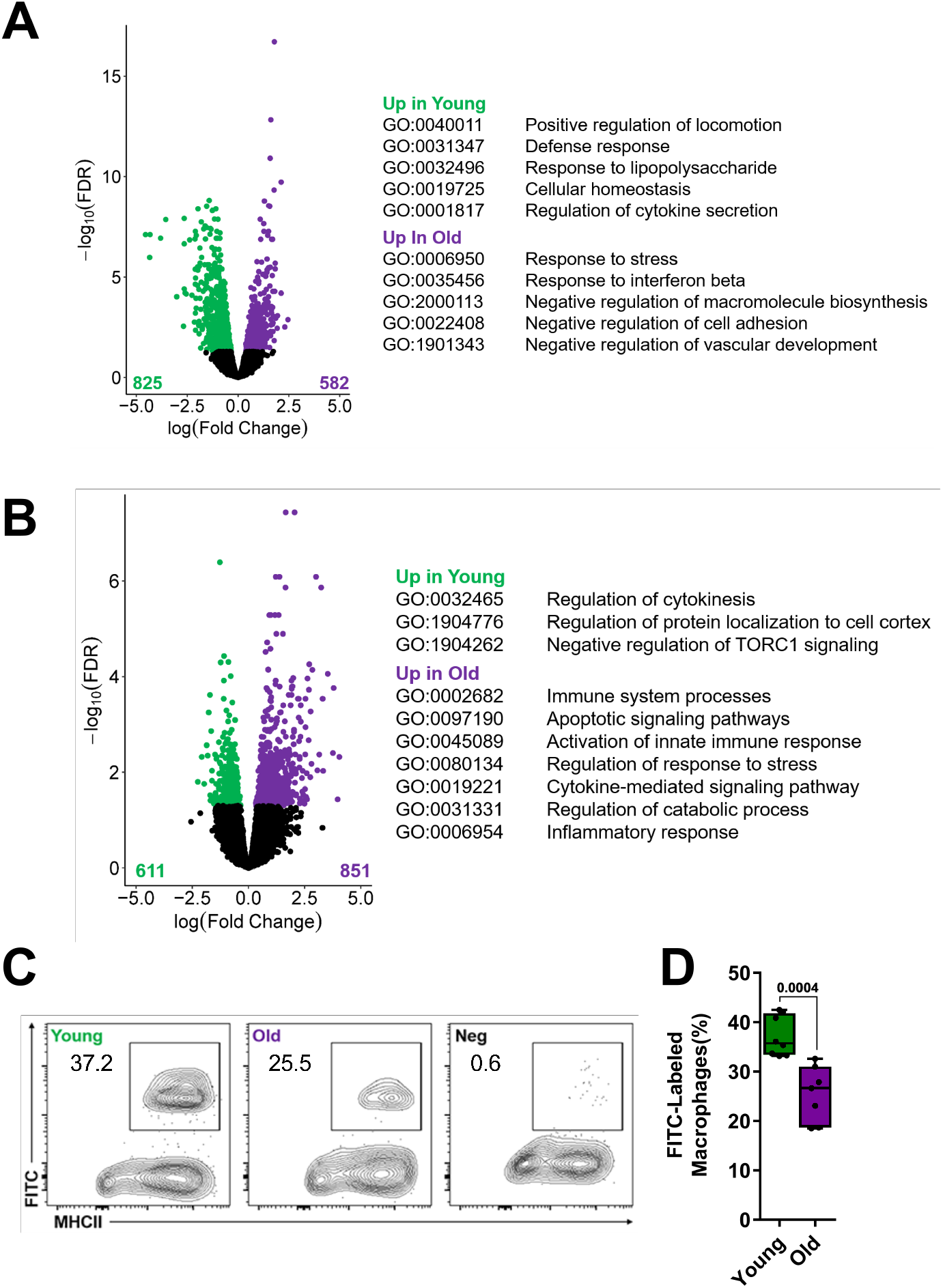
Tissue-resident macrophages from skeletal muscle from aged mice show impaired phagocytosis. (A) Skeletal muscle macrophages were flow cytometry sorted from naïve young (4 months old) and aged (20-24 months old) mice and analyzed using RNA-seq (n = 5 mice per group). Differentially-expressed genes (FDR q<0.05) between young (green dots) and aged (purple dots) mice are shown with representative genes and GO biological processes (See Tables S1-S2 for complete list of genes and GO processes). (B) Young (4 months old) and aged (20-24 months old) mice were infected with Influenza A virus (25 pfu for young mice, 15 pfu for aged mice) and harvested 60 days later. Skeletal muscle macrophages were flow cytometry sorted and analyzed using RNA-seq (n=5 mice per group). Differentially-expressed genes (FDR q<0.05) between young (green dots) and aged mice (purple dots) are shown with representative genes and GO biological processes (See Tables S3-S4 for complete list of genes and GO processes). (C) Single cell suspensions prepared from skeletal muscle of young (4 months old) and aged (20-24 months old) mice were incubated with antibodies to detect myeloid cells and serum opsonized Fluoresbrite microspheres. Uptake of the fluorescently labeled microspheres by macrophages was measured using flow cytometry. Non-opsonized beads were used as a negative control. Representative contour plots are shown, numbers indicate percentage from the parent population (gated on macrophages, see Figure 4A for gating strategy). (D) Refers to (C). Graph showing average percent of phagocytosed microspheres per macrophage (n=7-8 mice). A two-way ANOVA with Tukey post hoc corrections for comparison with more than three groups was used to show statistical differences. Box plot center lines are median, box limits are upper and lower quartiles, whiskers are minimal and maximal values, p values are shown on the graph. p values are shown on the graph. Each dot represents an individual animal.

### The phagocytic receptor *Mertk* is necessary for skeletal muscle repair after influenza A-induced pneumonia

MERTK is a member of the TAM family (Tyro3/Axl/Mer) tyrosine kinases that is expressed on macrophages, and is involved in recognizing and clearing apoptotic cells through phagocytosis (Lemke, 2013). We found that the level of mRNA encoding *Mertk* and its phosphatidylserine binding co-receptor *Gas6* were reduced in flow cytometry-sorted macrophages from aged compared with young mice (Figure 6A). Accordingly, we infected *Mertk^-/-^* and wild type mice with influenza A virus and measured skeletal muscle injury and recovery post-infection. Wild-type and *Mertk^-/-^*mice developed comparable levels of systemic IL-6 and Atrogin-1 expression in the skeletal muscle (Figure 6B, C), suggesting a similar level of muscle injury. However, measurements of voluntary distance run on a monitored exercise wheel showed that after an initial transient recovery, *Mertk^-/-^* mice failed to maintain their pre-infection levels of activity after pneumonia (Figure 6D, E). Additionally, we found increased accumulation of age-related lipofuscin granules in the muscle of young *Merkt^-/-^* mice compared to young wild type mice, similar accumulation of lipofuscin granules was observed in aged mice (Figure 6J, K and Figure S6). Like in *Cx3cr1* knockout mice, we observed increased numbers of skeletal muscle macrophages in *Mertk^-/-^* mice at baseline. However, the number of skeletal muscle macrophages in *Mertk^-/-^*mice did not increase during recovery from influenza A virus-induced pneumonia (Figure 6F) and there was no increase in the proportion of macrophages that were MHCII^low^ (Figure 6G), similar to aged mice. Consistent with these findings, *Mertk^-/-^*mice lacked expansion of MuSC and FAP during recovery from influenza A-induced pneumonia (Figure 6H, I). Thus, these data support a requirement for functional phagocytosis in recovery from influenza A pneumonia-induced skeletal muscle atrophy.

**Figure 6.**
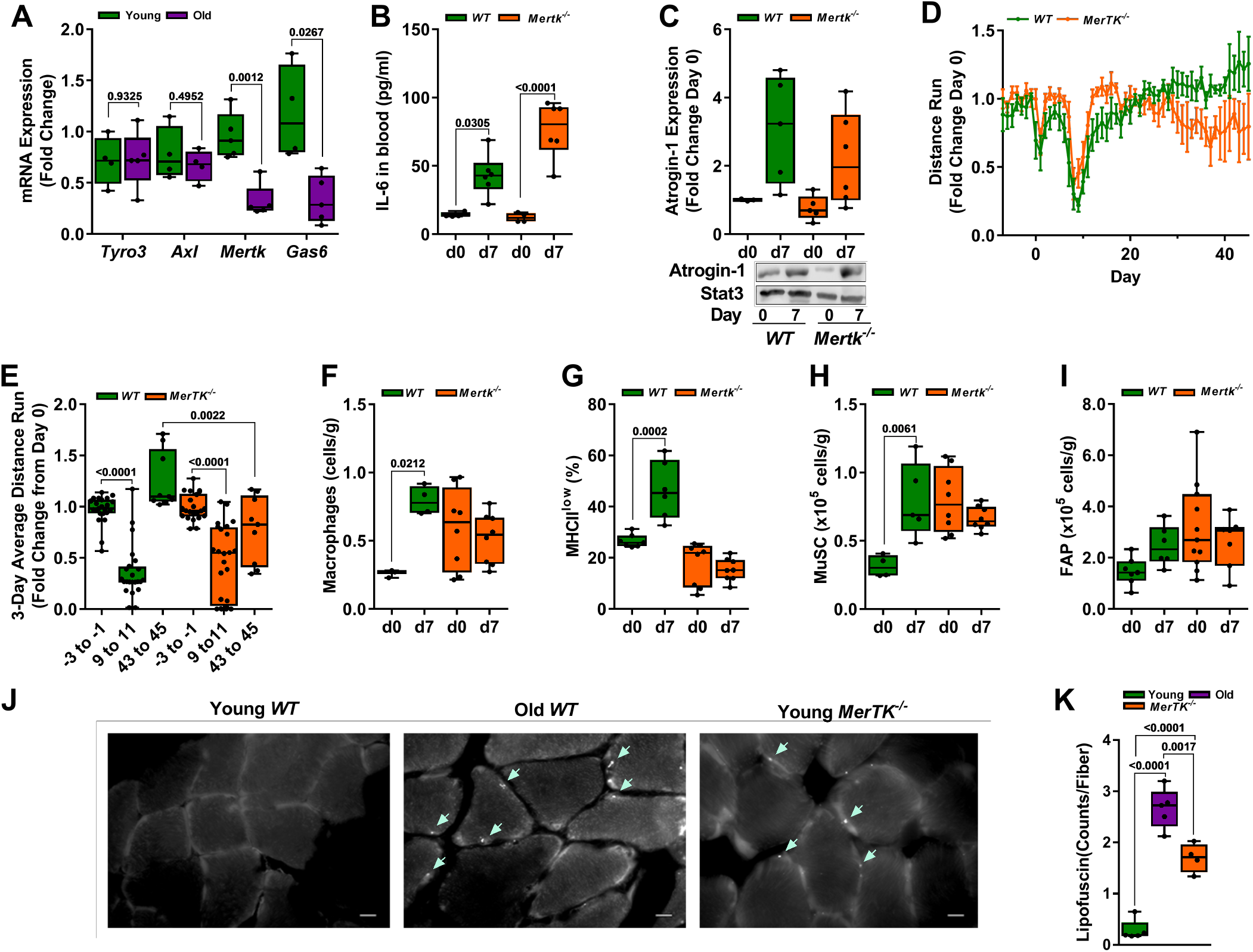
MERTK is necessary for recovery of skeletal muscle function after influenza A-induced pneumonia. (A) Skeletal muscle macrophages were isolated from naïve young (4 months: green bars) and aged (20-24 months: purple bars) mice and RNA was analyzed for TAM family members via real-time qPCR with SYBR green (n=5 mice). (B) Young (4 months old: orange bars) *Mertk^-/-^* mice and age-matched wild type control mice (green bars) were infected with influenza A virus (25 pfu) and harvested before the infection (Day 0) and 7 days after the infection. IL-6 was measured by ELISA in the serum (n=4-6 mice). (C) Young (4 months old: orange bars) *Mertk^-/-^* mice and age-matched wild type control mice (green bars) were infected with influenza A virus (25 pfu) and harvested before the infection (Day 0) and 7 days after the infection. Quantification (top panel) and representative Western blot (bottom panel) of Atrogin-1 expression in the mouse tibialis anterior muscle. GAPDH was used as a loading control (n=3-6 mice). (D) Prior to being infected with influenza A virus, young (4 months) *Mertk^-/-^*mice (orange lines) and age-matched wild type control mice (green lines) were exposed to voluntary wheel running for 14 days. They were then infected with 25 pfu of influenza A virus and voluntary wheel running was recorded for 50 days. The 8 pm to 6 am average wheel rotation/mouse, corrected to the initial 10-day average control is shown (n=15 mice per group). Graph shows distance run with means ± S.D. (E) Mice were treated as in (D). The 3-day average voluntary wheel running per cage is shown for day −3 to −1 prior to influenza A virus, day 9-11 post infection, and day 43 to 45 post infection (n=15 mice per group). (F) Young (4 months old, orange bars) *Mertk^-/-^* mice and age-matched wild type control mice (green bars) were infected with Influenza A virus (25 pfu) and harvested before the infection (Day 0) and 7 days after the infection. Skeletal muscle macrophages were enumerated using flow cytometry (n=4-8 mice). Values are expressed per gram of tissue. (G) Young (4 months old, orange bars) *Mertk^-/-^* mice and age-matched wild type control mice (green bars) were infected with Influenza A virus (25 pfu) and harvested before the infection (Day 0) and 7 days after the infection. MHCII^low^ skeletal muscle macrophages were enumerated using flow cytometry (n=4-8 mice). Values are expressed as a percent of total macrophages. (H) Young (4 months old, orange bars) *Mertk^-/-^* mice and age-matched wild type control mice (green bars) were infected with Influenza A virus (25 pfu) and harvested before the infection (Day 0) and 7 days after the infection. Skeletal muscle satellite cells (MuSC) were enumerated using flow cytometry (n=4-8 mice). Values are expressed per gram of tissue. (I) Young (4 months old, orange bars) *Mertk^-/-^* mice and age-matched wild type control mice (green bars) were infected with influenza A virus (25 pfu) and harvested before the infection (Day 0) and 7 days after the infection. Skeletal muscle fibroadipogenic progenitor cells (FAP) were enumerated using flow cytometry (n=4-8 mice). Values are expressed per gram of tissue. (J) Representative images of autofluorescent lipofuscin accumulation (identified by arrows) in unstained muscle tissue from naïve young (4 months old) mice, aged (20-24 months old) and young *Mertk^-/-^* (4 months old) mice. Bar is equal to 10 μm. (K) Graph depicting number of lipofuscin granules per fiber cross section counted manually in Image J. Five mice per group (an average of 3 fields of view per mouse). For all experiments, a two-way ANOVA with Tukey post hoc corrections for comparison with more than three groups was used to show statistical differences. Box plot center lines are median, box limits are upper and lower quartiles, whiskers are minimal and maximal values, p values are shown on the graph. Each dot represents an individual animal.

## DISCUSSION

Pneumonia is the most common infectious cause of death in the elderly, however, most elderly patients with access to modern healthcare survive pneumonia (Jain et al., 2015; Ortiz et al., 2013). In these survivors, skeletal muscle dysfunction is common and often persists for months or years after hospital discharge (Herridge et al., 2003). The elderly are at increased risk for the development of prolonged skeletal muscle dysfunction after pneumonia, however, the mechanisms that underlie this enhanced susceptibility have not been elucidated (Barreiro et al., 2015). In a murine model of influenza A pneumonia in which the dose of virus was titrated to result in similar mortality and weight loss, we found that young animals completely recovered skeletal muscle function after pneumonia, while aged animals did not. We used a combination of genetic lineage tracing, flow cytometry, transcriptomics and genetic loss of function approaches to show that a loss of phagocytic function in tissue-resident CX3CR1^+^skeletal muscle macrophages underlies the failure of skeletal muscle recovery in aged survivors of influenza A infection.

While a subpopulation of macrophages expressing the fractalkine receptor CX3CR1 in the skeletal muscle has been previously reported, nothing was known regarding the function or ontogeny of these cells (Arnold et al., 2015; Jin et al., 2018; Zhao et al., 2016). In young mice recovering from influenza A pneumonia, we observed expansion and downregulation of MHCII in skeletal muscle macrophages followed by muscle satellite cell proliferation. These findings were absent in mice deficient in *Cx3cr1*. Because macrophages are the only cell type in the skeletal muscle that expresses *Cx3cr1* (Schaum et al., 2018), these findings genetically link CX3CR1-expressing skeletal muscle macrophages with muscle satellite cells (MuSC) proliferation after influenza A-induced pneumonia. Moreover, we used a lineage tracing system and flow cytometry to show that, both in the steady state and after influenza A pneumonia, these skeletal muscle macrophages represent a *bona fide* tissue-resident macrophage population maintained in the skeletal muscle via local proliferation independent of recruitment of monocytes or monocyte-derived macrophages. Further, we found no evidence of infiltration by other myeloid cell populations or lymphoid populations in response to influenza A pneumonia. While the precise function of CX3CR1 in the skeletal muscle is not known, it may guide macrophages to areas of muscle loss induced by influenza A infection. For example, signaling through CX3CR1 guides monocytes and macrophages to areas of injury or inflammation, and during brain development, CX3CR1 is necessary to guide microglia to synapses where they are pruned from living neurons via a process referred to as “trogocytosis” (Paolicelli et al., 2011; Weinhard et al., 2018).

We used transcriptomic profiling to compare tissue-resident skeletal muscle macrophages in young and aged mice under naïve conditions and during recovery from influenza A infection. These data suggested the hypothesis that skeletal muscle macrophages from aged mice lose phagocytic function, which was confirmed by reduced uptake of serum-opsonized polystyrene beads in flow cytometry-sorted skeletal muscle macrophages from aged compared with young mice, and the accumulation of lipofuscin granules in skeletal muscle of aged and *Mertk^-/-^* mice. MERTK is a member of the TAM family of scavenger receptors (Mertk, Axl and Tyro3), which are activated during injury by interactions with Protein S or Gas6 bound to Annexin V on the surface of apoptotic cells (Lemke, 2013). Ligation of MERTK in macrophages results in the induction of phagocytosis through protein kinase C and Rac-mediated regulation of the cytoskeleton and the activation of AKT, leading to cellular proliferation and survival (Lemke and Rothlin, 2008). In addition, the activation of MERTK results in its association with the type I interferon receptor (IFNAR) to induce the Jak1/Stat1-mediated activation of suppressor of cytokine signaling 1 and 3 (SOCS1 and SOC3) (Lemke, 2013). These transcription factors inhibit inflammatory mediators and reduce expression of MHC class II molecules on the cell surface (Dalpke et al., 2001). We found that during recovery from influenza A infection, *Mertk^-/-^*mice largely phenocopied aged mice, in that they failed to expand tissue-resident macrophages, downregulate MHCII on tissue-resident macrophages or expand MuSC. The physiologic importance of these findings was reflected in their failure to recover normal exercise after influenza A infection, phenocopying aged mice. These findings strengthen the causal association between tissue-resident skeletal muscle macrophage function and muscle satellite cell proliferation. Furthermore, our findings are consistent with previous reports that MERTK is necessary for cardiac repair after myocardial infarction (DeBerge et al., 2017; Zhang et al., 2019).

Our findings fit into an emerging body of literature implicating impaired phagocytosis in tissue-resident macrophages with aging phenotypes. For example, Pluvinage performed a CRISPR screen to identify genes involved in microglia phagocytosis (Pluvinage et al., 2019). They found that CD22 was upregulated in aging microglia and impaired their phagocytic function. When they administered an antibody directed against CD22, cognitive function in aged mice improved. Future studies will answer whether improving phagocytic function in the tissue-resident macrophages in the skeletal muscle will be sufficient to revert influenza A pneumonia-induced muscle dysfunction in aged mice. Furthermore, our results support a role for tissue-resident macrophages in muscle progenitor cell proliferation. For example, co-culture of monocyte/macrophage populations with muscle progenitor cells induces their proliferation via the release of soluble factors in rodent and avian systems (Cantini and Carraro, 1995; Cantini et al., 2002; Chazaud et al., 2003; Merly et al., 1999; Saclier et al., 2013). In our model, however, we cannot determine whether impaired MuSC proliferation is related to the loss of paracrine signals induced in macrophages upon uptake of cellular debris or an inhibition of cellular proliferation by the debris that is not cleared by macrophages.

Voluntary wheel running in mice is an intuitive and compelling measure of the skeletal muscle dysfunction that impairs mobility in survivors of pneumonia. More importantly, our molecular findings are consistent with studies of muscle biopsies from patients with myopathy after critical illness. In these biopsies, investigators observed MuSC proliferation in the absence of inflammatory cell infiltration (Dos Santos et al., 2016). This differs from many murine models used to study muscle recovery, most of which involve direct injury to the muscle including trauma, freezing or cardiotoxin injection. In all of these injuries, inflammatory cells are recruited to the injured muscle including neutrophils, monocytes, monocyte-derived macrophages, regulatory T cells (Tregs) and other T cell populations, among others (Arnold et al., 2007; Heredia et al., 2013; Kuswanto et al., 2016; Teixeira et al., 2003; Varga et al., 2016; Zhang et al., 2014). Inflammatory cell recruitment in those models precluded the specific study of tissue-resident skeletal muscle macrophages. For example, two independent groups reported that deletion of monocyte-derived macrophages recruited in response to cardiotoxin-induced injury resulted in impaired satellite cell proliferation. In both of these studies, the strategies used to deplete monocyte-derived macrophages (CD11b-DTR and an antibody to M-CSF) would target both those monocyte-derived macrophages and the skeletal muscle tissue-resident macrophages we identified in this study (Arnold et al., 2007; Segawa et al., 2008). While Tregs have been reported to be important for skeletal muscle recovery after cardiotoxin injury (Kuswanto et al., 2016), we found that Treg number was reduced in both young and aged mice after influenza A infection. Nevertheless, Treg proliferation was higher in young compared with old mice and it is possible that factors released from resident tissue Tregs are important for recovery.

Our study has some limitations. First, we found that muscle loss after influenza A infection was sufficiently severe to induce myonuclear loss and the accretion of myonuclei in influenza A virus-infected mice (Mitchell and Pavlath, 2004) (Snijders et al., 2014) (McCarthy et al., 2011) (Egner et al., 2016), but we were unable to definitively determine the fate of the lost myonuclei in this model. Second, while *Mertk^-/-^* mice recapitulated the lack of cellular responses in aged animals, they transiently regained muscle function after injury. This may be explained by the high basal levels of MuSC in *Mertk^-/-^* relative to wild type mice, which allows short term, but unsustainable recovery of muscle function. Third, while we observed expansion of fibroadipogenic progenitors (FAP) during recovery from muscle injury in young, aged and *Mertk*-deficient mice, their role in muscle recovery is not certain. FAP cells have been described as important facilitators of MuSC behaviors, supporting their myogenic differentiation (Fiore et al., 2016), yet FAP cell persistence has been shown to have a detrimental effect on muscle tissue by contributing to fibrosis (Gonzalez et al., 2017) and ectopic fat accumulation (Uezumi et al., 2010). Thus, if tissue-resident macrophages contribute to FAP clearance, as has been reported for inflammatory, monocyte-derived macrophages (Lemos et al., 2015) some of the effects of aging or *Mertk* loss in macrophages might be attributed to the persistence of these cells. Finally, while the expression of *Cx3cr1* in the skeletal muscle is limited to macrophages, *Cx3cr1* is also expressed on monocytes and LYVE1-expressing macrophages associated with the vasculature and nerve fibers (Lim et al., 2018; Schaum et al., 2018). We therefore cannot completely exclude off target effects related to global deletion of *Cx3cr1* in our system.

The skeletal muscle represents approximately 40% of the total body mass (Janssen et al., 2000), and maintaining its function is critical for posture, breathing, locomotion, nutrient storage, and overall well-being. As a result, age-related sarcopenia is a major driver of frailty (Janssen et al., 2000; Topinkova, 2008). Our findings highlight the importance of impaired repair in the persistent, sometimes life-long loss of muscle function that develops in elderly survivors of pneumonia. Specifically, we suggest that a loss of phagocytic function in tissue-resident skeletal muscle macrophages during aging impairs their ability to promote muscle satellite cell proliferation necessary for skeletal muscle recovery. Given the prevalence of influenza A and other respiratory viral infections, it is likely that our findings have broad implications for elderly survivors of pneumonia.

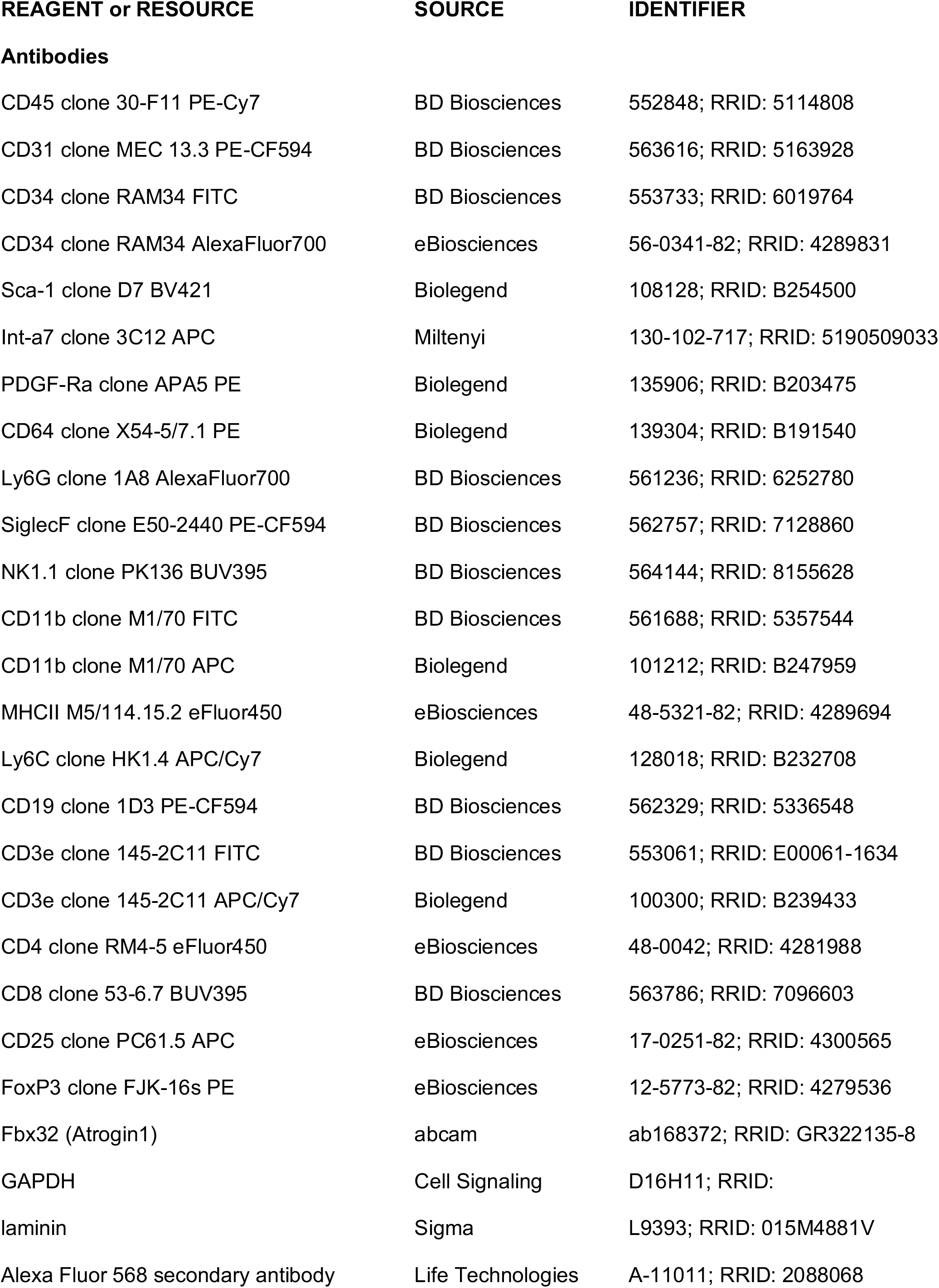

| REAGENT or RESOURCE | SOURCE | IDENTIFIER |
| --- | --- | --- |
| **Antibodies** |  |  |
| CD45 clone 30-F11 PE-Cy7 | BD Biosciences | 552848; RRID: 5114808 |
| CD31 clone MEC 13.3 PE-CF594 | BD Biosciences | 563616; RRID: 5163928 |
| CD34 clone RAM34 FITC | BD Biosciences | 553733; RRID: 6019764 |
| CD34 clone RAM34 AlexaFluor700 | eBiosciences | 56-0341-82; RRID: 4289831 |
| Sca-1 clone D7 BV421 | Biolegend | 108128; RRID: B254500 |
| Int-a7 clone 3C12 APC | Miltenyi | 130-102-717; RRID: 5190509033 |
| PDGF-Ra clone APA5 PE | Biolegend | 135906; RRID: B203475 |
| CD64 clone X54-5/7.1 PE | Biolegend | 139304; RRID: B191540 |
| Ly6G clone 1A8 AlexaFluor700 | BD Biosciences | 561236; RRID: 6252780 |
| SiglecF clone E50-2440 PE-CF594 | BD Biosciences | 562757; RRID: 7128860 |
| NK1.1 clone PK136 BUV395 | BD Biosciences | 564144; RRID: 8155628 |
| CD11b clone M1/70 FITC | BD Biosciences | 561688; RRID: 5357544 |
| CD11b clone M1/70 APC | Biolegend | 101212; RRID: B247959 |
| MHCII M5/114.15.2 eFluor450 | eBiosciences | 48-5321-82; RRID: 4289694 |
| Ly6C clone HK1.4 APC/Cy7 | Biolegend | 128018; RRID: B232708 |
| CD19 clone 1D3 PE-CF594 | BD Biosciences | 562329; RRID: 5336548 |
| CD3e clone 145-2C11 FITC | BD Biosciences | 553061; RRID: E00061-1634 |
| CD3e clone 145-2C11 APC/Cy7 | Biolegend | 100300; RRID: B239433 |
| CD4 clone RM4-5 eFluor450 | eBiosciences | 48-0042; RRID: 4281988 |
| CD8 clone 53-6.7 BUV395 | BD Biosciences | 563786; RRID: 7096603 |
| CD25 clone PC61.5 APC | eBiosciences | 17-0251-82; RRID: 4300565 |
| FoxP3 clone FJK-16s PE | eBiosciences | 12-5773-82; RRID: 4279536 |
| Fbx32 (Atrogin1) | abcam | ab168372; RRID: GR322135-8 |
| GAPDH | Cell Signaling | D16H11; RRID: |
| laminin | Sigma | L9393; RRID: 015M4881V |
| Alexa Fluor 568 secondary antibody | Life Technologies | A-11011; RRID: 2088068 |

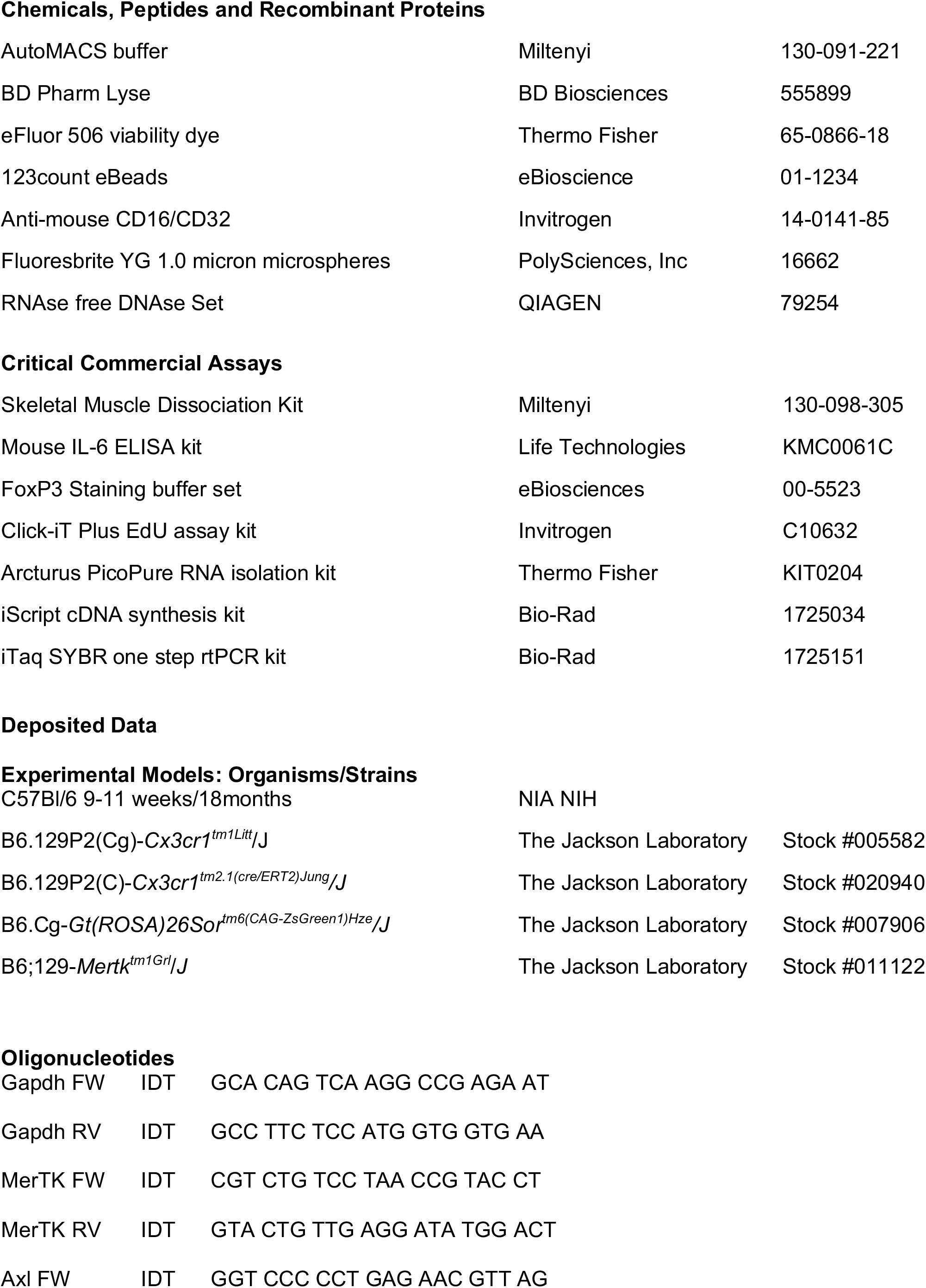

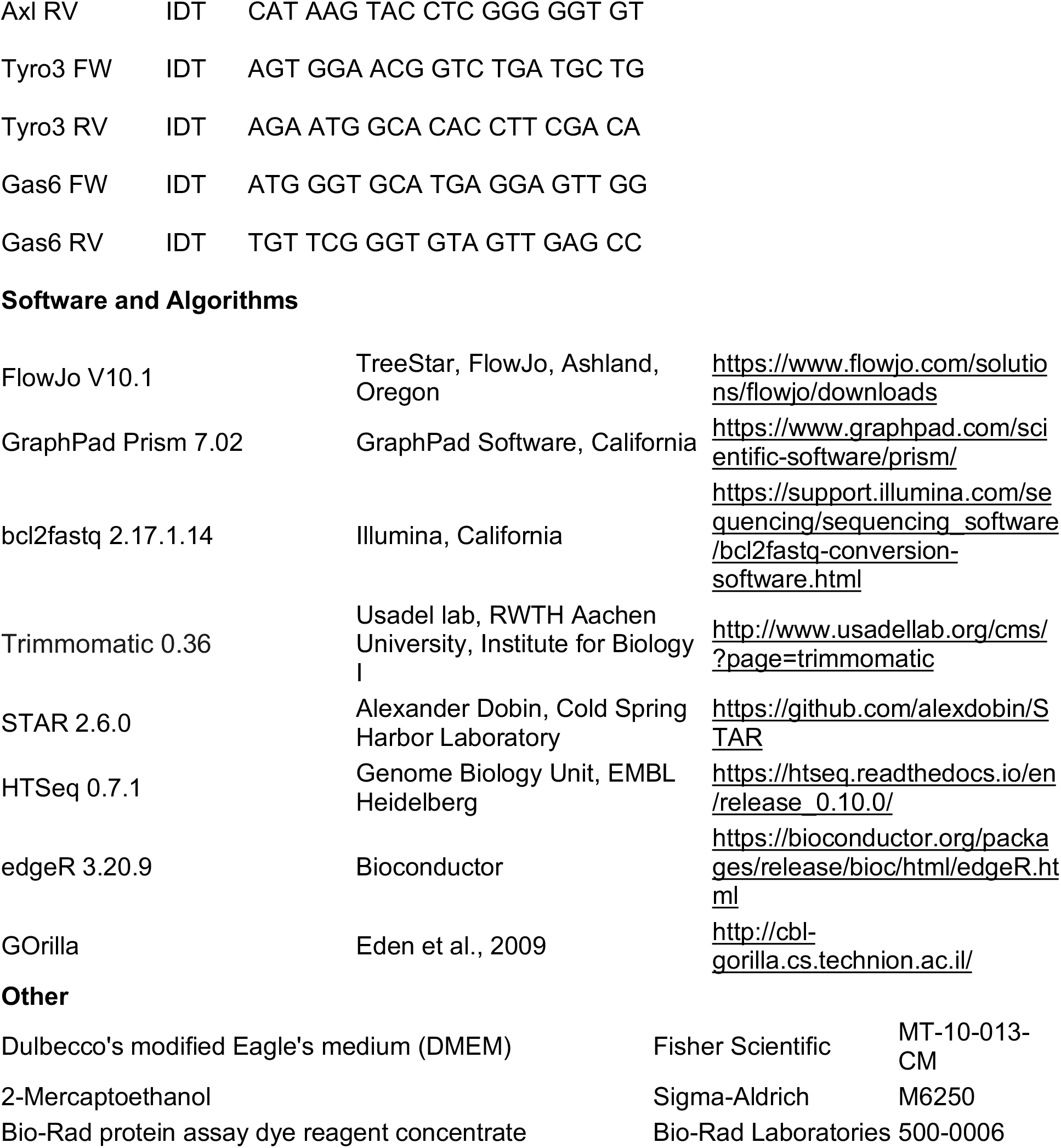

## CONTACT FOR REASGENT AND RESOURCE SHARING

Further information and requests for resources and reagents should be directed to and will be fulfilled by the Lead Contacts: Jacob Sznajder and Alexander Misharin.

## EPERIMENTAL MODEL AND SUBJECT DETAILS

### Mouse Models

All mouse procedures were approved by the Institutional Animal Care and Use Committee at Northwestern University (Chicago, IL, USA). All strains including wild-type mice are bred and housed at a barrier- and specific pathogen–free facility at the Center for Comparative Medicine at Northwestern University. All experiments were performed with littermate controls. Number of animals per group was determined based on our previous publications. Mice were housed at the Center for Comparative Medicine at Northwestern University, in microisolator cages, with standard 12 hr light/darkness cycle, ambient temperature 23°C and were provided standard rodent diet (Envigo/Teklad LM-485) and water *ad libitum*. Four and eighteen months-old C57BL/6J mice were obtained from Jackson laboratories or NIA colony. The C57BL/6J, *Cx3cr1^ER-Cre^*mice (Yona et al., 2013) and ZsGreen reporter (Madisen et al., 2010) mice were obtained from Jackson laboratories (Jax stocks 000664, 020940 and 007906, correspondingly) as were Cx3cr1*^gfp/gfp^* mice (Jax stock 005582).

## METHODS DETAIL

### Influenza-A infection

Murine-adapted, influenza A virus (A/WSN/33 [H1N1]) was provided by Robert Lamb, Ph.D., Sc.D., Northwestern University, Evanston, IL. Mice were anesthetized with isoflurane, their lungs were intubated orally with a 20-gauge Angiocath (Franklin Lanes, NJ), and then instilled with either sterile, phosphate-buffered saline (S) or mouse-adapted influenza A virus (IAV) as previously described (Radigan et al., 2019). Various doses in a range of 10-100pfu were tested to find an optimal sub-lethal dose matched to the age of the animal. Based on survival curves, we determined that 25 pfu was optimal for young animals, but a lower dose was required for aged animals, as a 25 pfu dose resulted in excessive mortality of old mice (Figure S1). We continuously observed mice infected with influenza A virus for signs of distress (slowed respiration, failure to respond to cage tapping, failure of grooming, huddling and fur ruffling). Mice that developed these symptoms were sacrificed and the death was recorded as an influenza A-induced mortality. Weight was measured prior to influenza infection and prior to harvest at 4 or 10 days post-infection, and recorded as percentage of weight loss from baseline (Radigan et al., 2019).

### Measurement of muscle dysfunction

Immediately prior to muscle harvest, forelimb skeletal muscle strength was assessed using a digital grip strength meter (Columbus Instruments, Columbus, OH) as described (Radigan et al., 2019). Grip strength was measured in each animal six successive times, and the average of the highest four values for each mouse was used. The same operator performed all tests. The mice were then terminally anesthetized with Euthasol (pentobarbital sodium/phenytoin sodium). The soleus and *extensor digitorum longus* (EDL) muscles were excised and tendon was trimmed under a microscope to assure optimal accuracy for weight measurement. The muscles were then blotted dry and weighed. Muscles were either frozen in liquid nitrogen-cooled isopentane for cryosectioning or snap-frozen in liquid nitrogen for protein extraction. Alternatively muscles were minced for flow cytometric analysis as described below.

### Voluntary Wheel Running

Mice, housed 3 per cage, were provided access to a trackable, low-profile saucer wheels (Med Associates, Inc., St. Albans, Vermont) 24 hours per day for 14-28 days to establish consistent running patterns. After establishing baseline, all mice were infected with IAV at an age-appropriate dose (Figure S1). Wheel rotations were gathered in one minute increments via wireless hub, and analyzed using wheel manager software (Med Associates, Inc., St. Albans, Vermont). Distances travelled per mouse per night (8 pm-6 am) were calculated for each night. Data were collected as the average of 3 mice per cage for 6 cages each, and running distance is expressed related to average nightly runs over 10 days pre-Influenza A.

### Immunohistochemistry and Fiber Size and Type Assessment

Soleus and *extensor digitorum longus* (EDL) serial transverse cryosections (8 μm) were obtained from the Northwestern University Mouse Histology and Phenotyping Laboratory and mounted on glass slides. Sections were fixed in 4% formaldehyde, permeabilized, and blocked. Immunostaining was performed with laminin primary antibody (1:50 dilution; Sigma; Catalog: L9393) followed by Alexa Fluor 568-conjugated secondary antibody (1:200 dilution; Life Technologies; Catalog: A-11011). Images were acquired with a Zeiss LSM 510 confocal microscope using a 40x objective (Northwestern University Center for Advanced Microscopy) and analyzed using Zeiss LSM5 Image Browser software. Fiber size was studied by measuring the fibers’ minimal inner diameter (at least 100 fibers per muscle), defined as the minimum diameter from inner border to inner border, passing through the center of the muscle fiber. This parameter has been shown to be very insensitive to deviations from the “optimal” cross-sectioning profile, as compared with direct measure of fiber cross-sectional area (Radigan et al., 2019). Cross-sectional area (CSA) was calculated using this diameter, and results were expressed as mean CSA ± S.E.

### Bronchoalveolar lavage

Saline fluid (0.8 ml) was lavaged then aspirated from the lungs of mice through tracheal angiocatheter. Cell counts were measured by Trypan blue exclusion and the automated Countess system. To measure the cell differential, 200 μl of BAL fluid was placed in Cytospin funnels and spun 2,000 rpm 5 min onto a glass slide. Cells were stained by Wright-Giemsa and manually counted and identified.

### IL-6 measurements

IL-6 expression assays were performed using a kit from Life technologies (KMC0061C) according to the manufacturer’s instructions. Assays were performed on either BAL fluids (acquired as described above) or from 50 μl/well blood plasma drawn from posterior vena cava using a 23-gauge needle, centrifuged to remove cells, and stored at −80°C. Analysis for each mouse was done in triplicate, and a minimum of 5 mice were analyzed/condition.

### Flow Cytometry and Cell Sorting

Mice were fully-perfused through the right ventricle with 20 ml of PBS, and hind-limb muscle was removed. Tissue was cut into small pieces with scissors, transferred into C-tubes (Miltenyi, Auburn, CA) and processed with a Skeletal Muscle Dissociation kit according to manufactures instructions (Miltenyi, Auburn CA). To achieve a single-cell suspension, 2 rounds of dissociation were performed using a GentleMACS dissociator (Miltenyi), followed by 3 rounds of filtration through Falcon 100-μm, 70-μm then 40-μm nylon mesh filter units (Thermo Fisher #352340, 352350 and 352360 respectively) into polypropylene 50ml Falcon tubes, followed by centrifugation at 1300rpm in an Eppendorf 5810R centrifuge for 15 minutes. Any remaining red blood cells were lysed by resuspending the pellet in 1ml/tube BD Pharm Lyse (BD Biosciences, San Jose, CA), and were transferred to flow tubes, pelleted by low speed centrifugation 5 minutes, and resuspended in 0.5ml/tube HBSS containing 0.5 μl/tube viability dye eFluor 506 (eBioscience, San Diego, CA). Cells were stained in viability dye for 15 minutes in the dark at room temperature, followed by 2X washes in 1ml MACS buffer (Miltenyi, Auburn, CA). After live/dead cell staining, pellets are resuspended in 150 μl/tube FcBlock Anti-mouse CD16/CD32 (Invitrogen) for 30 minutes in the dark at 4°C to block non-specific binding. After blocking, cells were divided to three separate tubes at 50 μl each, then stained with a mixture of fluorochrome-conjugated antibodies in 50 μl/tube, and incubated for an additional 30 minutes at 4°C in the dark. After antibody incubation, the cells were washed 2X in MACS buffer, and then fixed in 2% paraformaldehyde for 15 minutes at room temperature in the dark, washed 2X in HBSS, resuspended in 200 μl HBSS, then stored at 4°C overnight. Data were acquired on a BD LSR Fortessa II flow cytometer using BD FACSDiva software, and compensation and data analyses were performed using FlowJo software (TreeStar, Ashland, OR). Prior to cytometric analysis, 50 μl 123count eBeads (Invitrogen) were added to 200 μl each sample to allow accurate cell counts/gm of tissue. “Fluorescence minus one” controls were used when necessary. Cell populations were identified using sequential gating strategy (Figure 3A, 4A and 5A). For Foxp3 detection, a staining kit for intracellular antigens was used (eBiosciences, San Diego, CA). Cells were fixed and permeabilized according to directions, before incubation with Foxp3 antibody prior to analysis. Cell sorting was performed at Northwestern University RLHCCC Flow Cytometry core facility on SORP FACSAria III instrument (BD) with the same optical configuration as LSR II, using a 100-µm nozzle and 40 psi pressure.

To measure proliferation, EdU was diluted in sterile saline from 100 mg/ml stock in DMSO, and injected intraperitoneally at 2 mg/mouse, 16 hours prior to tissue harvest (Misharin et al., 2017). Single-cell suspensions were divided in thirds, and stained with antibody panels as described in (Figure 3A, 4A and 5A) with FITC replaced in each panel. After surface antibody staining cells are fixed and undergo a Click-IT EdU detection reaction using an Alexa-Flour 488 picolyl azide, Click-IT Plus EdU Flow cytometry assay kit (Invitrogen).

### Quantitative RT-PCR

Ten thousand skeletal muscle macrophages were sorted on a FACSAria 6-Laser Sorter, into 200 μl PicoPure lysis buffer and stored at −80°C. RNA was harvested following manufactures directions for Arcturus PicoPure RNA isolation kit (Applied Biosystems, Foster City, CA). Total RNA was quantified using a NanoDrop 2000c (Thermo Scientific). cDNA was generated using iScript cDNA synthesis kit (Bio-Rad, Hercules CA) in a Bio-Rad T100 thermal cycler, followed by qPCR with SYBRgreen (Bio-Rad, Hercules CA) and a Bio-Rad CFX Connect real-time system. The following primers were used:

Gapdh (F-GCA CAG TCA AGG CCG AGA AT, R-GCC TTC TCC ATG GTG GTG AA) MerTK (F-CGT CTG TCC TAA CCG TAC CT, R-GTA CTG TTG AGG ATA TGG ACT) Axl (F-GGT CCC CCT GAG AAC GTT AG, R-CAT AAG TAC CTC GGG GGT GT) Tyro3 (F-AGT GGA ACG GTC TGA TGC TG, R-AGA ATG GCA CAC CTT CGA CA) Gas6 (F-ATG GGT GCA TGA GGA GTT GG, R-TGT TCG GGT GTA GTT GAG CC)

### Western Blot Analysis

Quadriceps muscle tissue was homogenized with a Tissue Tearor (BioSpec Products, Inc) for 1 minute in ice-cold lysis buffer (Nonidet P-40 1%, glycerol 10%, NaCl 137 mM, Tris-HCl pH 7.5 20 mM) containing protease (1 Complete Mini, EDTA-free tablet, Roche) and phosphatase (sodium fluoride 30 mM, β-glycerophospate 250 mM, sodium orthovanadate 1mM) inhibitors. Samples were centrifuged at 15,000 rpm for 10 minutes at 4°C and the supernatant was collected. Protein concentrations were determined with Protein Assay Dye (Bio-Rad) in a 96 well plate measured against BSA protein standard dilutions at 595 nm with a Bio-Rad iMark microplate reader (Bio-Rad, Hercules CA). Equal amount of protein (40-80 total μg) were loaded on a 9% SDS-PAGE gel, and Western blot analysis was performed as described previously (Radigan et al., 2019). Incubation with primary antibodies was performed overnight at 4°C. Immunoblots were quantified by densitometry using Image J 1.46r (National Institutes of Health, Bethesda, MD) or Image Studio Software (LI-COR Inc., Lincoln, NE). The following antibodies were used: Rabbit Monoclonal to Fbx32 (Abcam; catalog: ab168372; 1:1000) or Rabbit monoclonal to GAPDH (D16H11) (Cell Signaling; catalog: 5174s; 1:1000).

### Transcriptome Profiling via mRNA-Seq

Ten thousand skeletal muscle macrophages were sorted on a FACSARIA 6-Laser Sorter, into 200 μl PicoPure lysis buffer and stored at −80°C. RNA was harvested following manufactures directions for Arcturus PicoPure RNA isolation kit (Applied Biosystems, Foster City, CA). RNA quality was assessed with the 4200 TapeStation System (Agilent). Samples with an RNA integrity number (RIN) lower than 7 were discarded. RNA-seq libraries were prepared from 0.5 ng total RNA using the SMARTer Stranded Total RNA-Seq Kit v2 - Pico Input Mammalian kit. Libraries were quantified and assessed using the Qubit Fluorimeter (Invitrogen) and 4200 TapeStation. Libraries were sequenced on NextSeq 500 instrument (Illumina) at 75 bp length, single end reads. The average reading depth across all experiments exceeded 6×10^6^ per sample and over 94% of reads had a Q-score greater than 30. For RNA-seq analysis, reads were demultiplexed using bcl2fastq (version 2.17.1.14). Read quality was assessed with FastQC. Low quality baseballs were trimmed using trimmomatic (version 0.36). Reads were then aligned to the *Mus musculus* reference genome (with mm10 assembly) using the STAR aligner (version 2.6.0). Counts tables were generated using HTSeq (version 0.6.1). Raw counts were processed in R (version 3.4.4) using edgeR (version 3.20.9) to generate normalized counts. Negative binomial likelihood with default setting followed by generalized linear models fitting was used to estimate differentially expressed genes. FDR q-values were used to correct for multiple comparisons and a value of 0.05 was used as a threshold to indicate statistical significance (see online code). Gene Ontology analysis was performed using GOrilla on two unranked gene lists. The RNA-seq datasets are available at GEO: GSEXXXXXX.

### *In Vitro* Phagocytosis

After enzymatic digestion 50 μl of the single cell suspensions were incubated with 35 μl of myeloid antibody cocktail with 15 μl/tube Fluoresbrite YG 1.0 micron microspheres (Polysciences Inc., Warrington, PA) that had been opsonized for 30 minutes at 37°C in by mixing 1 drop beads with 1ml 50% FBS/HBSS, followed by 2X wash/resuspension in HBSS 1 ml (non-opsonized beads were used as a negative control). Cell suspensions were allowed to uptake beads for 30 minutes at 37°C in the dark with shaking, followed by 2 washes in HBSS to remove any unbound beads, and fixation in 2% paraformaldehyde, followed by flow cytometry with a FITC negative myeloid cocktail to detect bead uptake in myeloid cell populations.

## QUANTIFICATION AND STATISTICAL ANALYSIS

### Statistical Analysis

Differences between groups were determined according to ANOVA. When ANOVA indicated a significant difference, individual differences were examined using *t* tests with a Tukey correction for multiple comparisons, as indicated. All analyses were performed using GraphPad Prism, version 5.00 (GraphPad Software, San Diego CA). Data are shown as means ± SEMs. *P* < 0.05 was considered statistically significant in all tests.

**Supplementary table S1:** List of differentially expressed genes between tissue-resident skeletal muscle macrophages from naïve young and old mice.

**Supplementary table S2:** Gene ontologies for the differentially expressed genes between tissue-resident skeletal muscle macrophages from naïve young and old mice.

**Supplementary table S3:** List of differentially expressed genes between tissue-resident skeletal muscle macrophages from young and old mice, 60 days after influenza A infection.

**Supplementary table S4:** Gene ontologies for the differentially expressed genes between tissue-resident skeletal muscle macrophages from young and old mice, 60 days after influenza A infection.

## Acknowledgments

Northwestern University Flow Cytometry Facility, Center For Advanced Microscopy, and Pathology Core Facility are supported by NCI Cancer Center Support Grant P30 CA060553 awarded to the Robert H. Lurie Comprehensive Cancer Center. This research was supported in part through the computational resources and staff contributions provided by the Genomics Computing Cluster (Genomic Nodes on Quest) which is jointly supported by the Feinberg School of Medicine, the Center for Genetic Medicine, and Feinberg’s Department of Biochemistry and Molecular Genetics, the Office of the Provost, the Office for Research, and Northwestern Information Technology.

## Funding

Satoshi Watanabe is supported by MSD Life Science Foundation, Public Interest Incorporated Foundation, Japan, and David W. Cugell and Christina Enroth-Cugell Fellowship Program at Northwestern University. GR Scott Budinger is supported by NIH grants ES013995, HL071643, AG049665, The Veterans Administration Grant BX000201. Jacob I Sznajder is supported by NIH grants HL147070, HL071643 and AG049665. Alexander V Misharin is supported by NIH grants HL135124, AG049665, AI135964.

The authors declare no competing financial interests.

## Author contributions

Constance Runyan: designed the study, performed experiments, analyzed results, wrote manuscript. Lynn C. Welch: performed experiments, analyzed results, wrote manuscript. Emilia Lecuona, Masahiko Shigemura, Luciano Amarelle, Raul Piseaux-Aillon, Kinola J.N. Williams, Nikita Joshi, Kiwon Nam, Alexandra C. McQuattie-Pimentel, Yuliya Politanska, Ziyan Lu, Lango Sichizya, Satoshi Watanabe: performed experiments, analyzed results. Hiam Abdala-Valencia: designed the study, performed experiments, analyzed results. Nikolay Markov: performed bioinformatics analysis, wrote manuscript. GR Scott Budinger: designed and supervised the study, wrote manuscript, provided reagents, resources and funding. Jacob I Sznajder and Alexander V. Misharin: designed and supervised the study, performed analysis, wrote manuscript, provided funding for the project. All authors discussed the results and commented on the manuscript.

---

**Figure S1.**
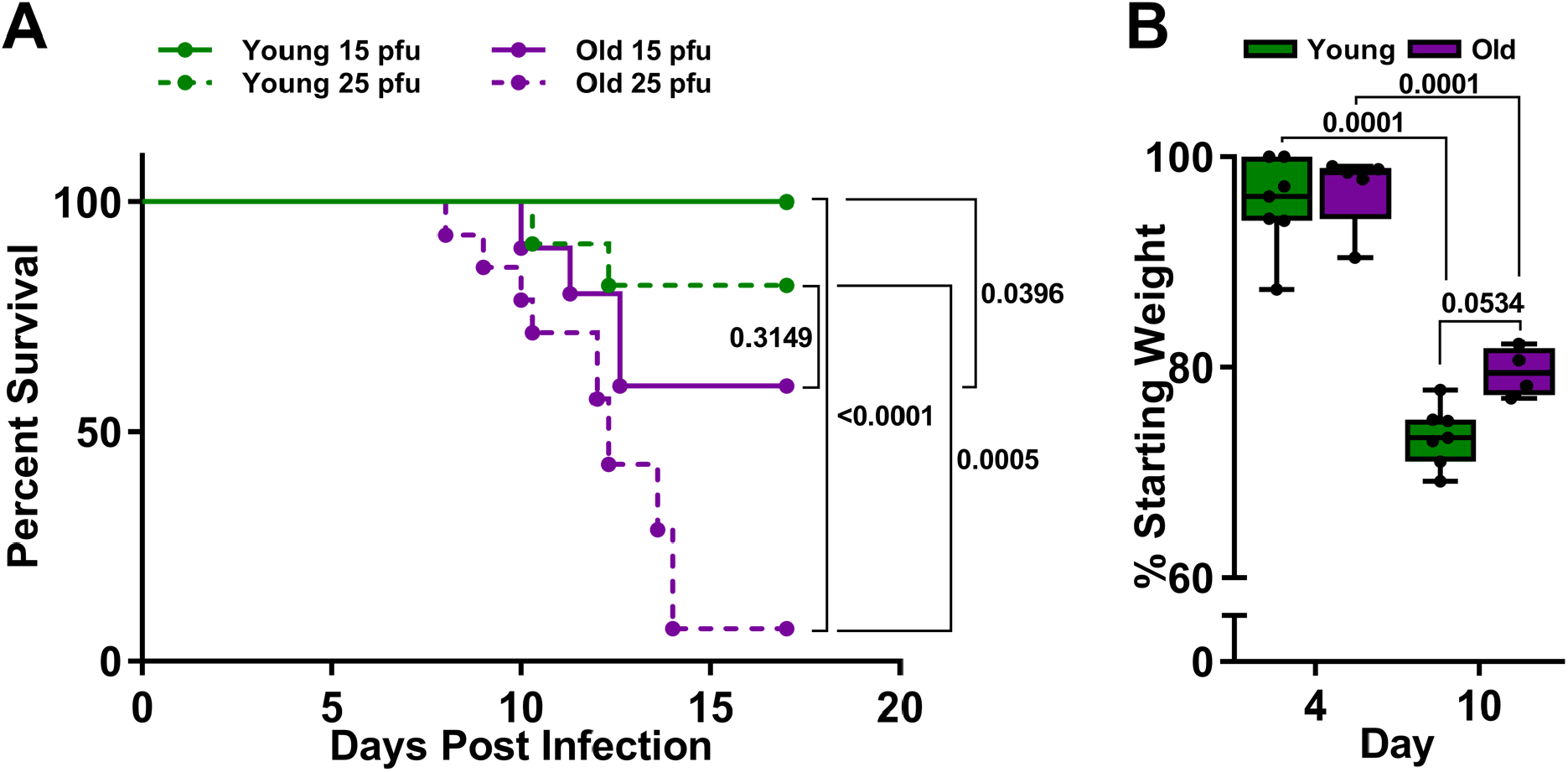
Influenza A virus causes disproportionate mortality in old mice. Refers to Figure 1. (A) Young (4 months old; green lines) and old (20-24 months old; purple lines) mice were infected with either 15 pfu (solid lines) or 25 pfu (dotted lines) of influenza A virus (A/WSN/33) (IAV). A Kaplan-Meier survival curve is shown (n = 9-14 mice per group). Log-rank (Mantel-Cox) tests were performed and p values are shown on graph. Each dot represents a single animal. (B) Young (4 months old; green bars) mice were infected with 25 pfu of IAV and old (20-24 months old; purple bars) mice were infected with 15 pfu influenza A virus (IAV). Percentage of starting total body weight is shown for day-4 and day-10 post-IAV infection (n=5-7 mice per group). Box plot center lines are median, box limits are upper and lower quartiles, whiskers are minimal and maximal values, a two-way ANOVA with Tukey post-hoc corrections for comparison with more than three groups was used to determine statistical differences, and p values are shown on graph. Each dot represents a single animal.

**Figure S2.**
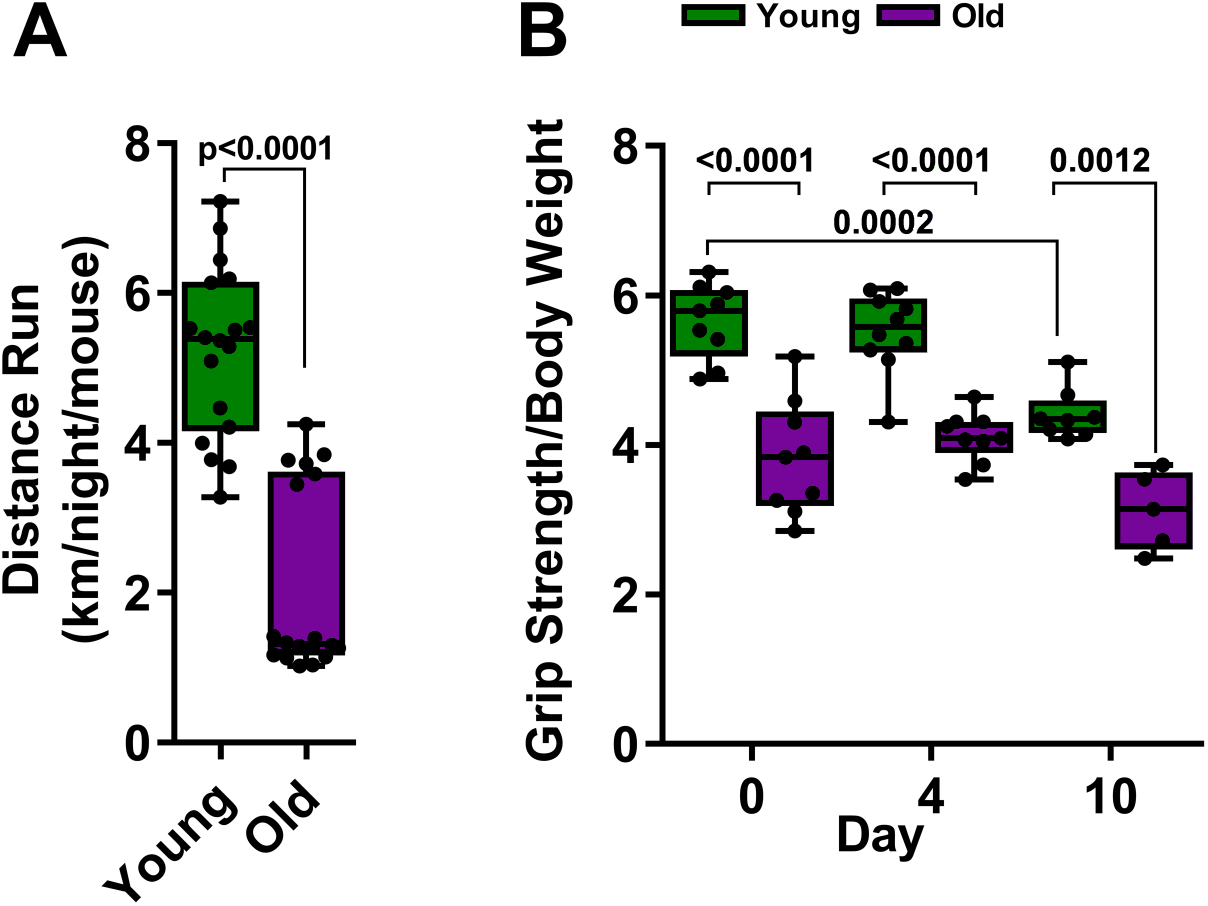
Old mice fail to recover skeletal muscle function after influenza A infection (refers to Figure 1). (A) Young (4 months old; green bars) and old (20-24 months old; purple bars) mice were allowed voluntary access to a running wheel in their home cage. Average running distance from 8 pm-6 am was collected over 10 consecutive nights (n=12 mice per group). Box plot center lines are median, box limits are upper and lower quartiles, whiskers are minimal and maximal values, unpaired t test, and p values are shown on graph. Each dot represents a single animal. (B) Young (4 months old; green bars) and old (20-24 months old; purple bars) mice were infected with IAV (25 pfu for young mice, 15 pfu for old mice). Forelimb grip strength was measured (n=4-10 mice per group). Box plot center lines are median, box limits are upper and lower quartiles, whiskers are minimal and maximal values, a two-way ANOVA with Tukey post-hoc corrections for comparison with more than three groups was used to determine statistical differences, and p values are shown on graph. Each dot represents a single animal.

**Figure S3.**
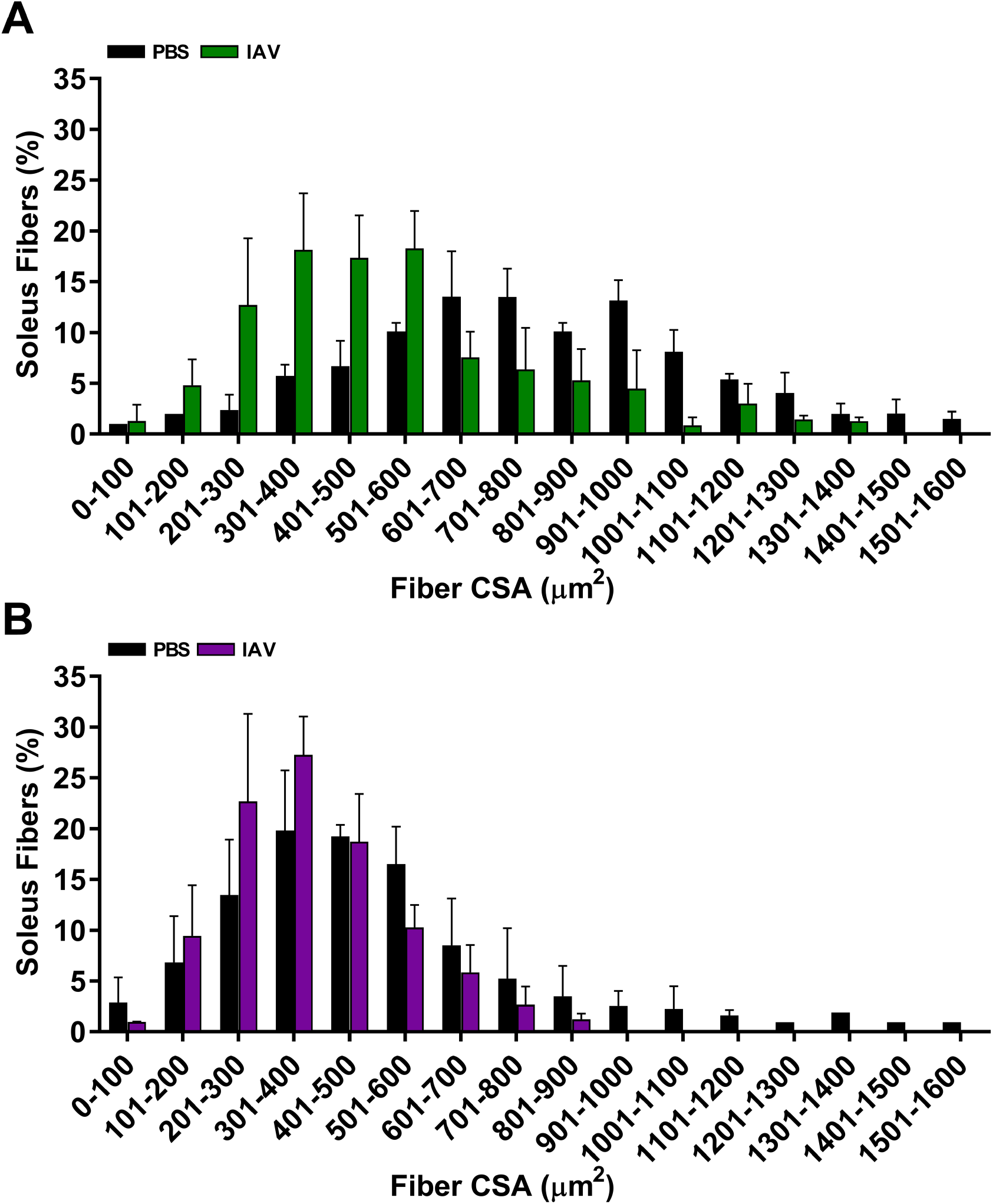
Aged mice have lower skeletal muscle fiber diameter at baseline. Young mice lose more muscle fiber cross sectional area after influenza A infection compared with old mice. (A) Young mice (4 months old) were infected with 25 pfu influenza A virus (IAV; green bars), mice instilled with PBS were used as a control (black bars). Mice were harvested for analysis 10 days later. Soleus muscles were excised, frozen and cryosectioned. Soleus cross-section area was determined using cross-sections immunostained for laminin. A histogram depicting the fiber size distribution in young mice is shown (n = 3 mice (naïve), 5 mice after influenza A infection). Bars represent means ± S.D. (B) Aged mice (20-24 months old) were infected with 15 pfu influenza A virus (IAV; purple bars). Mice instilled with PBS were used as a control (black bars). Mice were harvested for analysis 10 days later. Soleus muscles were excised, frozen and cryosectioned. Soleus cross-section areas were determined using cross-sections immunostained for laminin. A histogram depicting the fiber size distribution in old mice is shown (n = 4 mice (naïve), 5 mice after influenza A infection). Bars represent means ± S.D.

**Figure S4.**
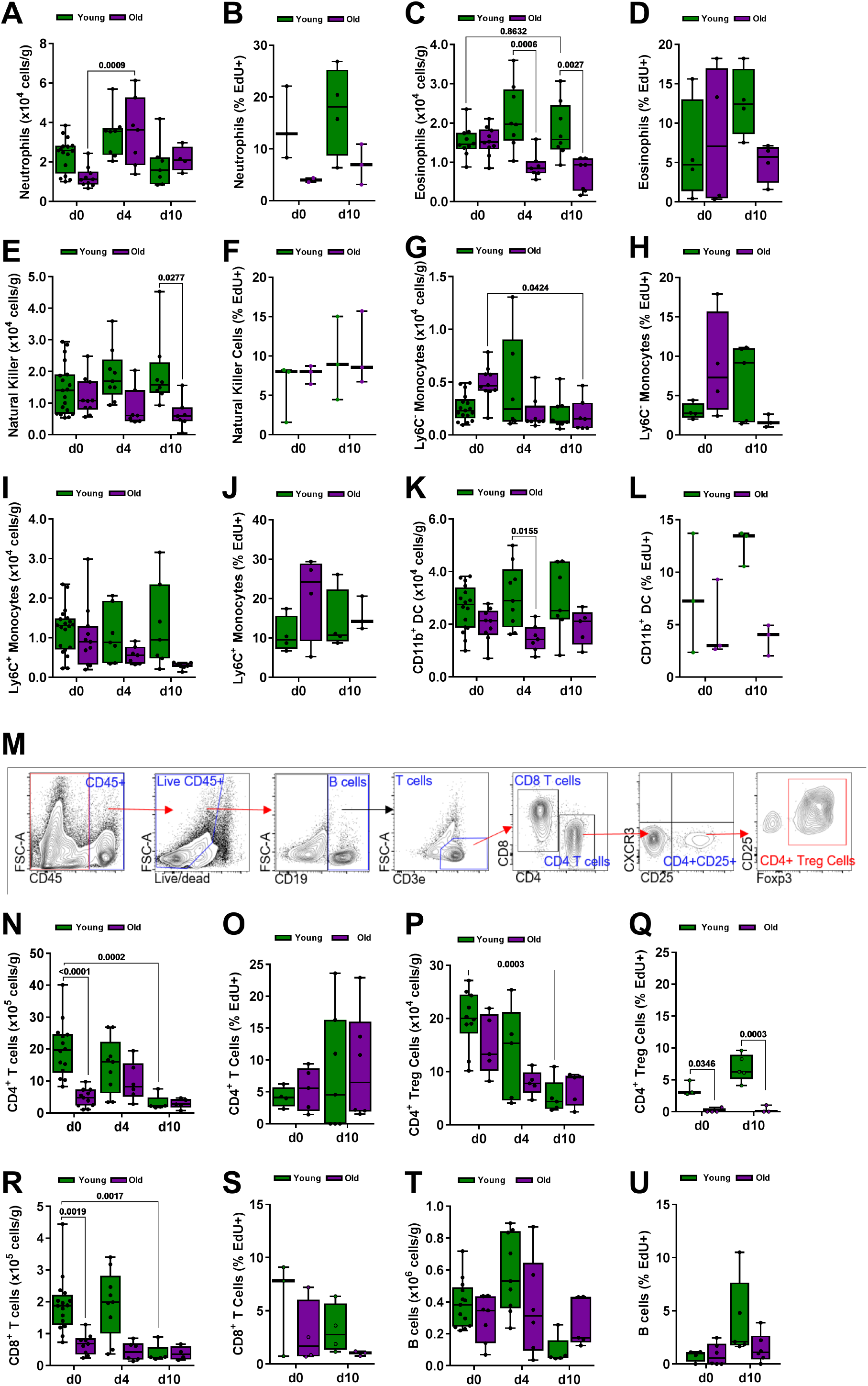
Influenza A pneumonia does not result in the recruitment of myeloid or lymphoid cell populations to the skeletal muscle. Refers to Figure 4. For all experiments, young (4 months old; green bars) and aged (20-24 months old; purple bars) mice were infected with influenza A virus (25 pfu for young mice, 15 pfu for aged mice) and analyzed before infection (Day 0) and 4 and/or 10 days after infection. A two-way ANOVA with Tukey post hoc corrections for comparison with more than three groups was used to show statistical differences, p values are shown on the graph. Box plot center lines are median, box limits are upper and lower quartiles, whiskers are minimal and maximal values, p values are shown on the graph. Each dot represents an individual animal. (A) Graph depicting total number of neutrophils. Values are expressed per gram of tissue (n=5-16 mice). (B) Graph representing EdU^+^ neutrophils. Values are expressed as a percentage of total neutrophils (n=3-4 mice). (C) Graph depicting total number of eosinophils. Values are expressed per gram of tissue (n=7-10 mice). (D) Graph representing EdU^+^ eosinophils. Values are expressed as a percentage of total eosinophils (n=3-4 mice). (E) Graph depicting total number of natural killer cells. Values are expressed per gram of tissue (n=7-19 mice). (F) Graph representing EdU^+^ natural killer cells. Values are expressed as a percentage of total natural killer cells (n=3-4 mice). (G) Graph depicting total number of Ly6C^-^ monocytes. Values are expressed per gram of tissue (n=6-16 mice). (H) Graph representing EdU^+^ Ly6C^-^ monocytes. Values are expressed as a percentage of total Ly6C^-^ monocytes (n=3-4 mice). (I) Graph depicting total number of Ly6C^+^ monocytes. Values are expressed per gram of tissue (7-19 mice). (J) Graph representing EdU^+^ Ly6C^+^ monocytes. Values are expressed as a percentage of total Ly6C^+^ monocytes (n=3-4 mice). (K) Graph depicting total number of CD11b^+^ dendritic cells. Values are expressed per gram of tissue (n=7-16 mice). (L) Graph representing EdU^+^ dendritic cells. Values are expressed as a percentage of total dendritic cells (n=3-4 mice). (M) Gating strategy used for flow cytometric analysis of B and T cells. (N) Graph shows number of CD4 T cells in skeletal muscle per gram of tissue before infection (Day 0) and 4 and 10 days after influenza A infection (n=5-11 mice). (O) Graph shows percentage of EdU^+^ CD4 T cells, of total CD4 T cells, in skeletal muscle before infection (Day 0) and 10 days after influenza A infection (n=4-6 mice). (P) Graph shows number of CD4 Treg cells in skeletal muscle per gram of tissue before infection (Day 0) and 4 and 10 days after influenza A infection (n=5-11 mice). (Q) Graph shows percentage of EdU^+^ CD4 Treg cells, of total CD4 Treg cells, in skeletal muscle before infection (Day 0) and 10 days after influenza A infection (n=4-6 mice). (R) Graph shows number of CD8 T cells in skeletal muscle per gram of tissue before infection (Day 0) and 4 and 10 days after influenza A infection (n=5-11 mice). (S) Graph shows percentage of EdU^+^ CD8 T cells, of total CD8 T cells, in skeletal muscle before infection (Day 0) and 10 days after influenza A infection (n=4-6 mice). (T) Graph shows number of B cells in skeletal muscle per gram of tissue before infection (Day 0) and 4 and 10 days after influenza A infection (n=5-11 mice). (U) Graph shows percentage of EdU^+^ B cells, of total B cells, in skeletal muscle before infection (Day 0) and 10 days after influenza A infection (n=4-6 mice).

**Figure S5.**
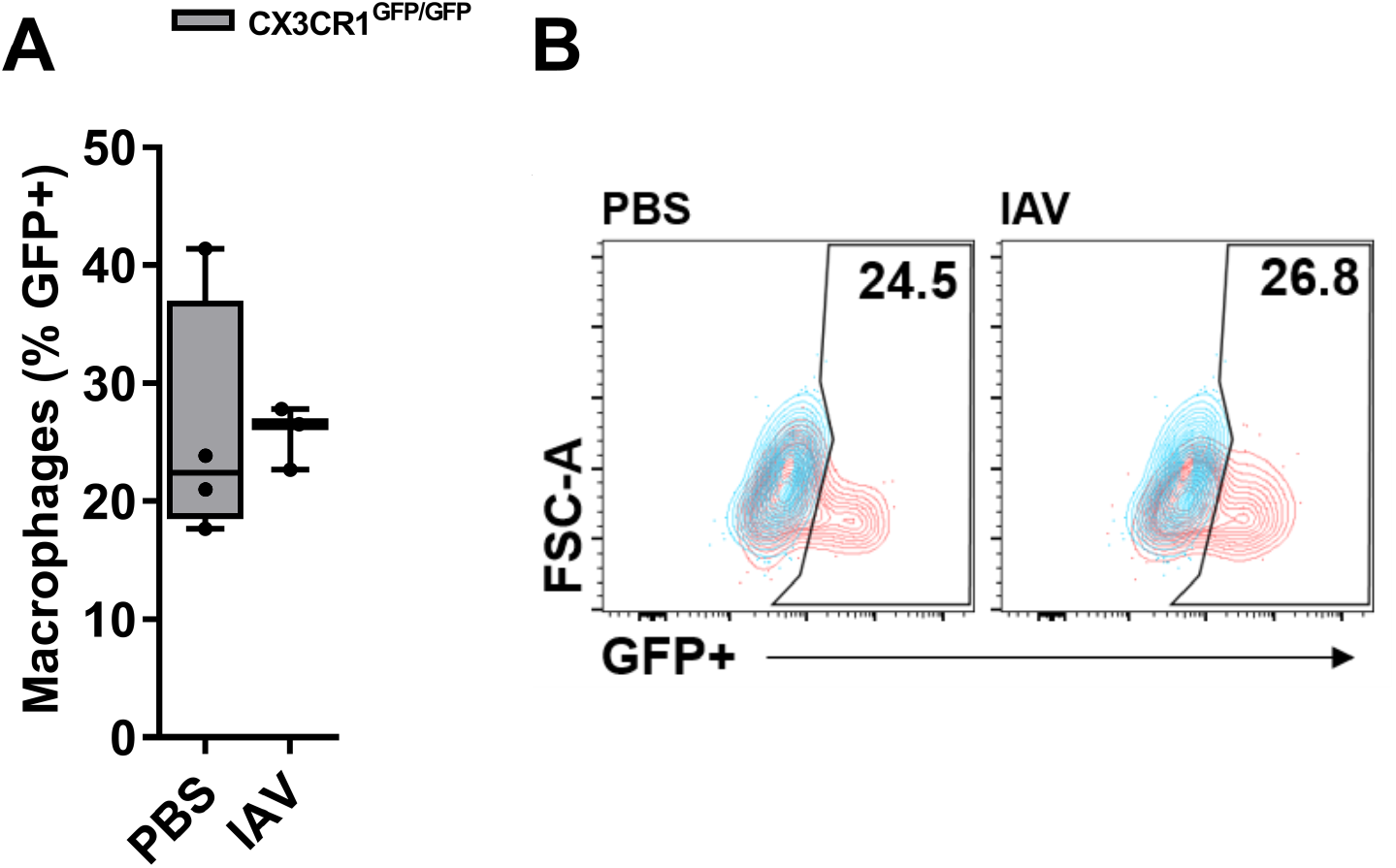
Identification of a CX3CR1^+^ population of tissue resident skeletal muscle macrophages. (A) Heterozygous mice in which GFP is knocked in to the *Cx3cr1* gene (*B6.129P2(Cg)-Cx3cr1^tm1Litt^/J* (Jackson labs 005582)), referred to in the text as *Cx3cr1^Gfp/+^* were infected with the influenza A virus (25pfu) or sham infected with PBS and harvested 10 days later for analysis using flow cytometry. (B) Representative flow cytometry panels show GFP^+^ (CX3CR1^+^) macrophages in red, overlaid over WT mouse macrophages in blue. Numbers represent percent of cells expressing GFP.

**Figure S6.**
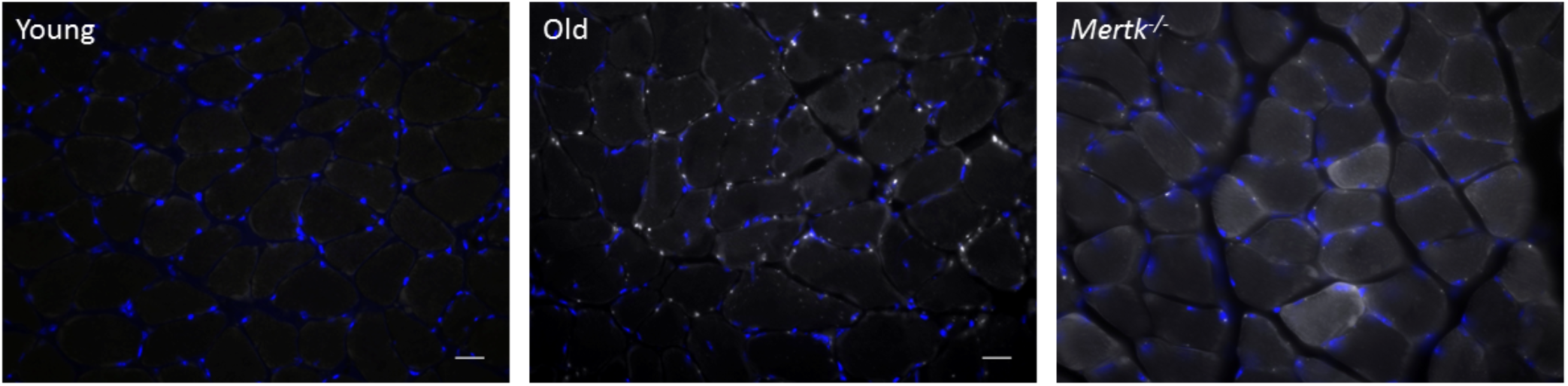
Impaired phagocytosis results in increased lipofuscin accumulation. Soleus muscles were harvested from naïve young mice (4 months), old mice (20-24 months) or young *Mertk^-/-^* mice (4 months), and stained for DAPI (blue). Autofluorescent lipofuscin granules (white). Scale bar=40 μm,

